# The ChickenGTEx pilot analysis: a reference of regulatory variants across 28 chicken tissues

**DOI:** 10.1101/2023.06.27.546670

**Authors:** Dailu Guan, Zhonghao Bai, Xiaoning Zhu, Conghao Zhong, Yali Hou, The ChickenGTEx Consortium, Fangren Lan, Shuqi Diao, Yuelin Yao, Bingru Zhao, Di Zhu, Xiaochang Li, Zhangyuan Pan, Yahui Gao, Yuzhe Wang, Dong Zou, Ruizhen Wang, Tianyi Xu, Congjiao Sun, Hongwei Yin, Jinyan Teng, Zhiting Xu, Qing Lin, Shourong Shi, Dan Shao, Fabien Degalez, Sandrine Lagarrigue, Ying Wang, Mingshan Wang, Minsheng Peng, Dominique Rocha, Mathieu Charles, Jacqueline Smith, Kellie Watson, Albert Johannes Buitenhuis, Goutam Sahana, Mogens Sandø Lund, Wesley Warren, Laurent Frantz, Greger Larson, Susan J. Lamont, Wei Si, Xin Zhao, Bingjie Li, Haihan Zhang, Chenglong Luo, Dingming Shu, Hao Qu, Wei Luo, Zhenhui Li, Qinghua Nie, Xiquan Zhang, Zhe Zhang, Zhang Zhang, George E. Liu, Hans Cheng, Ning Yang, Xiaoxiang Hu, Huaijun Zhou, Lingzhao Fang

## Abstract

Chicken is a valuable model for understanding fundamental biology, vertebrate evolution and diseases, as well as a major source of nutrient-dense and lean-protein-enriched food globally. Although it is the first non-mammalian amniote genome to be sequenced, the chicken genome still lacks a systematic characterization of functional impacts of genetic variants. Here, through integrating 7,015 RNA-Seq and 2,869 whole-genome sequence data, the Chicken Genotype-Tissue Expression (ChickenGTEx) project presents the pilot reference of regulatory variants in 28 chicken tissue transcriptomes, including millions of regulatory effects on primary expression (including protein-coding genes, lncRNA and exon) and post-transcriptional modifications (alternative splicing and 3’ untranslated region alternative polyadenylation). We explored the tissue-sharing and context-specificity of these regulatory variants, their underlying molecular mechanisms of action, and their utility in interpreting adaptation and genome-wide associations of 108 chicken complex traits. Finally, we illustrated shared and lineage-specific features of gene regulation between chickens and mammals, and demonstrated how the ChickenGTEx resource can further assist with translating genetic findings across species.

**One-Sentence Summary:** The ChickenGTEx provides a multi-tissue reference of regulatory variants for chicken genetics and genomics, functional genomics, precision breeding, veterinary medicine, vertebrate evolution and even human biomedicine.

## Introduction

The chicken (*Gallus gallus domesticus*) is not only a globally significant source of protein-rich food through both meat and egg production, but also a fundamental model species. In 2021, the farming industry achieved a staggering production of 111 million tons of eggs and 137 million tons of poultry meat worldwide (https://www.fao.org). Due to its distinct phylogenetic placement as well as its genetic and physiological characteristics, the chicken is also served as a well-recognized model organism in both fundamental and applied research (*1*, *2*), studies of domestication, genome editing, system biology, virology, immunology, oncology, and evolution (*3–7*). The chicken retains a remarkable range of phenotypic variation in terms of morphology, physiology, and behavior, primarily driven by artificial selection and breed specialization. Such extensive variation for a wide range of features is ideal for investigating the genetic architecture underlying complex traits. One example of such traits is dwarfism, which is characterized by a short stature and is observed in various forms in chickens, including sex-linked dwarfism, autosomal dwarfism, and the bantam phenotype, according to their physiological and genetic properties (*8*, *9*). In addition, long-term bidirectional selection lines have been established in chickens to study how polygenetic selection influences complex traits such as body weight (*10*) and feather pecking (*11*).

The Red Jungle Fowl (*G. gallus*), the ancestor of domestic chicken, was one of the first food-producing animals that had its genome assembled (*1*). Recently, a near complete version of the chicken reference genome assembly was reported, revealling distinct sequence and epigenetic features of microchromosomes (*12*). Several population-scale studies of chicken genome variation have focused on various aspects of its evolution, including speciation and domestication (*13–15*), signatures of selection(*14*, *16*, *17*), admixture and introgression(*14*, *18*, *19*), feralization(*20*, *21*), and phenotypic adaptation(*22–24*). Meanwhile, linkage mapping and genome-wide association studies (GWAS) have identified tens of thousands of genomic loci associated with numerous complex traits in chickens (*5*, *25–27*). As most genetic variants behind such adaptive evolutionary and complex traits are non-coding, a systematic annotation of regulatory variants in the chicken genome becomes indispensable for understanding their underlying genetic regulatory circuitry (*28–30*). The expression quantitative trait locus (eQTL) analysis is presently the most powerful approach to measure regulatory effects of sequence variants on individual gene expression in their native genomic and cellular contexts (*31*), as documented in the human Genotype-Tissue Expression (GTEx) project series of studies (*32–34*) and eQTL Catalogue in humans (*35*). In contrast, previous eQTL studies in chickens have been limited in sample size, the number of studied genomic features, and tissue/cell types (*36–40*). For instance, in an intercross population of 125 chickens, Johnsson et al. (2015) identified 6,318 *cis*- eQTL that influence female femoral gene expression, as measured by microarrays (*36*).

To fully unlock the genetic code of the chicken genome, the Chicken Genotype-Tissue Expression (ChickenGTEx) project, as part of the international Farm animal GTEx (FarmGTEx) initiative (*41*), has been lunched to build a comprehensive reference panel of regulatory variants based on the chicken transcriptome in various biological contexts (e.g., development, sex and environmental exposure). In this pilot study, through analyzing 7,015 bulk RNA-Seq datasets from 52 tissues/cell types (hereafter referred to as “tissues”) and 2,869 whole genome sequences (WGS) from over 100 breeds/lines (hereafter referred to as “breeds”) worldwide, we systematically associated approximately1.5 million genomic variants with five transcriptomic phenotypes in 28 chicken tissues with sufficient sample size (ranging from 44 to 741). We then explored tissue-sharing and context-dependent patterns of these regulatory variants, their underlying molecular mechanisms of action, and their utility in deciphering GWAS loci of 108 complex traits *via* multiple complementary integrative methods such as transcriptome-wide association studies (TWAS), colocalization, and Mendelian Randomization (MR). Additionally, we compared gene regulation and the phenotypic implications between chicken and three mammalian species (*i.e.* human, cattle and pig). Altogether, our study provides novel and profound insights into the regulatory hierarchy of genetic variation in chicken transcriptomes and complex phenotypes, providing the first large-scale mapping of regulatory variants in the chicken genome and their links to complex phenotypes. Meanwhile, the atlas of regulatory variants identified in this study will facilitate the genetic improvement of chicken populations worldwide in health, production, and resilience and inform a wide range of genetic and genomic research in animal and plant species. Furthermore, we have also well-developed a ChickenGTEx online resource that is freely accessible at http://chicken.farmgtex.org.

## Results

### Harmonizing large transcriptome and genome datasets in chickens

We analyzed 8,668 bulk RNA-Seq samples using a uniform pipeline, yielding 304.4 billion clean reads. After filtering out low-quality data, 7,015 samples remained for subsequent analyses, representing 28 tissues (**fig. S1, Tables S1** and **2**). Based on gene expressions, samples were clustered well regarding their tissue types (**fig. 1a, fig. S2**). Across all the tissues, an average of 23,056 (94.7% of all annotated genes) genes were expressed (Transcripts per Million, TPM > 0.1) (**Table S3**), showing patterns of ubiquitous or tissue-specific expression (**fig. S3**). An average of 1,938 tissue-specific genes were then detected across tissues (**fig. S3d**), and their functions recapitulated the known tissue biology (**fig. 1b**, **fig. S3e**, **Table S4, URL**). For instance, a total of 1,425 genes were specifically and highly expressed in the bursa of Fabricius, a bird-specific primary lymphoid organ, which were significantly enriched in the immune response to bacteria (**Table S4**). An average of 54.7% of tissue-specific genes could be linked to at least one tissue-specific promoter/enhancer (**fig. S4a-d**). For instance, *MSLNL* with bursa- specific promoters and enhancers showed a specific expression in the bursa (**fig. S4d**). An average of 114 genes exhibited sex-biased expression across 18 tissues (FDR < 0.01), among which 17 genes were shared in all these tissues and located in sex chromosomes (**figs. S4e-f, Table S5**). This was in agreement with the notion of incomplete sex-chromosome dosage compensation in chickens(*42*). In addition, out of 45 genes associated with Mendelian traits in chickens(*43*), 41 showed tissue-specific expression (**fig. S5a**). For instance, *SLCO1B3* is the causal gene of blue eggshell in chickens(*44*), which was specifically and highly expressed in the liver and had liver-specific promoters and enhancers (**fig. S5b**).

**Fig. 1.**
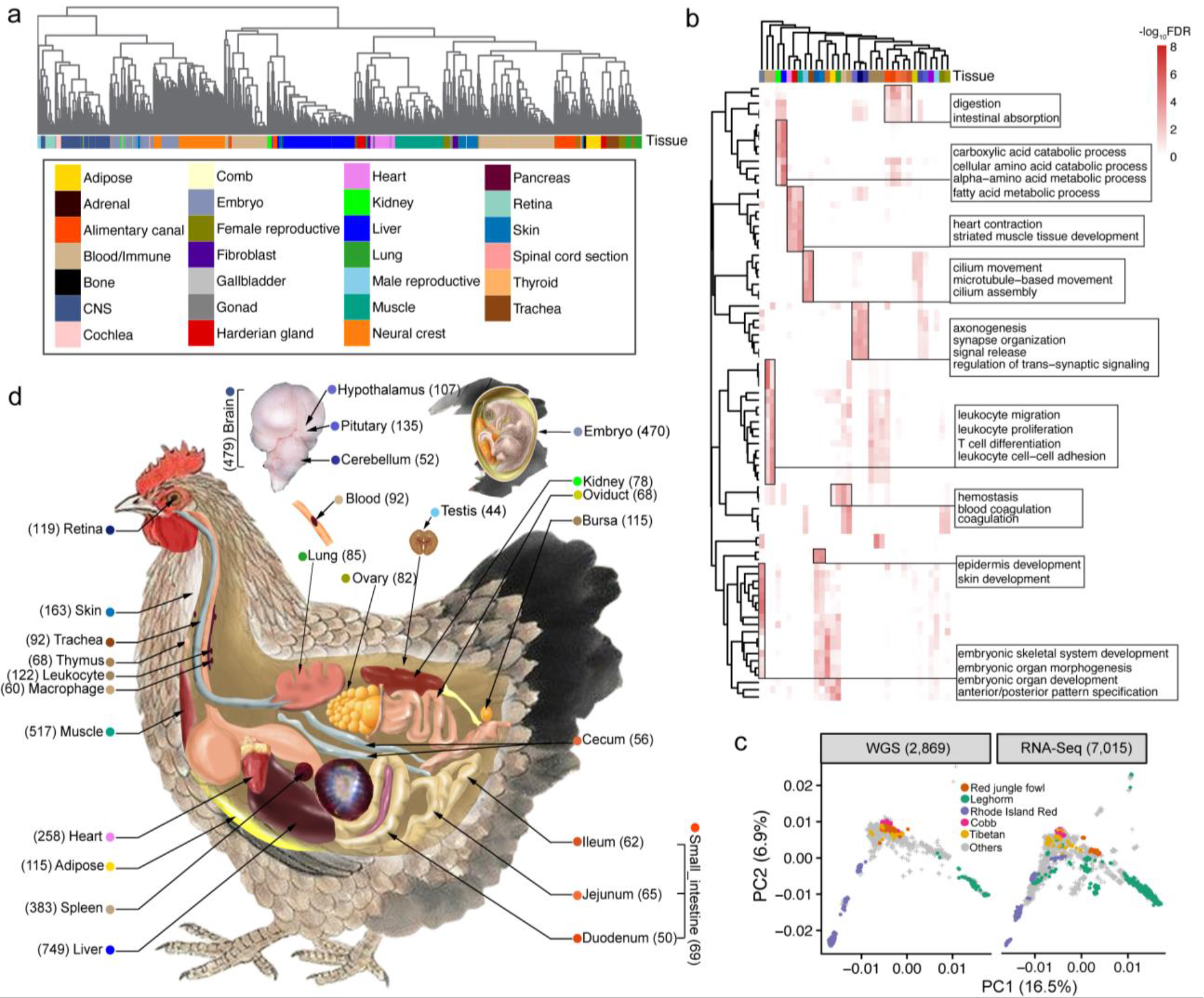
Data summary in the pilot phase of ChickenGTEx. (**a**) Hierarchical clustering of 7,015 RNA-Seq samples. Distance between samples was calculated using 1-*r*, where *r* is the Pearson correlation coefficient calculated from gene expression values (quantified as Transcripts per Million, TPM) of 5,000 genes with the highest expression variance (measured by standard deviation) across samples. (**b**) Functional enrichment of tissue-specific genes based on the Gene Ontology (GO) database. The color scale from light to deep means a negative logarithm of false discovery rate (FDR) at the base of 10, obtained by the clusterProfiler 4.0 package with default settings(*103*). (**c**) Scatterplots depicting principal component analysis (PCA) of 2,869 whole-genome sequence (WGS, left) and 7,015 RNA-Seq samples (right). PCA was carried out using 1.52 million SNP genotypes shared by both WGS and RNA-Seq datasets. (**d**) Illustration of tissue types used in molecular quantitative trait loci (molQTL) mapping. Sample sizes (in the bracket) and colors of all the 28 tissues with sample sizes over 40 are depicted.

To further annotate the function of chicken genes with this extensive transcriptome data, we conducted gene co-expression network and transcript assembly analyses. Based on co-expression analysis, we identified 3,583 co-expression modules containing 25,023 genes, 41.3% (10,332) of which were not functionally annotated in the current Gene Ontology (GO) database (**figs. S6**). In the set of 2,940 unannotated protein-coding genes, 56.3% (1,654) were able to be assigned to co-expression modules. Compared to annotated genes, these unannotated genes exhibited more tissue-specificity, lower gene expression level, and small proportion of chicken-human orthologous genes (**fig. S6d**). For instance, 8 unannotated genes were co-expressed with 12 annotated genes in the muscle, which were significantly enriched in myeloid cell development and erythrocyte differentiation networks (**fig. S6f)**. Through the transcript assembly analysis, we predicted 247,383 transcripts at 48,800 loci, including 184,374 protein-coding transcripts derived from 17,215 loci, 13,140 lncRNA transcripts from 3,436 loci, and 49,869 other noncoding RNA transcripts from 34,350 loci (**fig. S7**, URL). Of all these predicted transcripts, 90% were not annotated in the previous reference and 4-10% were even transcribed from novel genomic loci (**fig. S7d** and **g**). For instance, we observed a new transcript on chromosome 2 was highly and specifically expressed in the testis (**fig. S7h**).

To obtain genotypes of these RNA-Seq samples, we called ∼9 million single nucleotide polymorphisms (SNPs) from bulk RNA-Seq data using the GATK best practice pipeline(*45*) (**fig. S8**). To impute missing genotypes, we built a chicken multi-breed genotype imputation reference panel consisting of 2,869 global WGS data sets, which had a similar population composition as the RNA-Seq data (**fig. 1c, Table S6**). Adopting a missing rate of 0.6, the imputation accuracy of 1.5 million SNPs reached 97% (**figs. S8b-h**). The independent datasets from three different chicken breeds confirmed a high concordance rate (> 90%) between imputed genotypes and those directly called from WGS data of the same individuals (**fig. S8g**). After removing duplicated samples based on their genetic relatedness, 28 tissues (each consisting of over 40 individuals) were retained for subsequent molecular quantitative trait loci (molQTL) mapping (**fig. 1d**).

### Discovery of molQTL

To comprehensively explore the genetic regulation of the chicken transcriptome, we conducted *cis*-molQTL mapping for five molecular phenotypes, including protein-coding gene expression (eQTL), lncRNA expression (lncQTL), exon expression (exQTL), splicing variation (sQTL), and 3’UTR alternative polyadenylation (3a’QTL), across 28 chicken tissues (**fig. 2a, fig. S2, figs. S9- 10)**. In total, 13,983 (92.9%) of 15,046 tested protein-coding genes, 11,685 (74.3%) of 15,720 lncRNAs, 124,423 (76.0%) of 163,812 exons, 9,669 (61.5%) of 15,405 loci with alternative splicing events, and 8,798 (74.1%) of 11,880 loci with 3’UTR alternative polyadenylation (3’UTR APA) were significantly (FDR < 0.05) regulated by at least one genetic variant in at least one tissue, and are thus referred to as eGenes, lncGenes, exGenes, sGenes and 3a’Genes, respectively. All the molQTL tended to be enriched around transcription start sites (TSS) and transcription end sites (TES), while 3a’QTL and sQTL showed a higher enrichment in TES and gene body, respectively, compared to other molQTL (**fig. 2b**, **figs. S11a-e**). Furthermore, an average of 73.6% (10,288) of eGenes, 40.5% (3,914) of sGenes, 60.7% (75,527) of exGenes, 58.9% (6,886) of lncGenes, and 7.3% (640) of 3a’Genes were regulated by more than one independent variant (eVariants) across tissues (**fig. 2c, fig. S12**). The further fine-mapping analysis for molQTL with SuSiE (*46*) revealed 2,887, 2,366, 2,053, 12,409 and 1,572 potential causal variants for eGenes, sGenes, lncGenes, exGenes and 3a’Genes, respectively (URL).

**Fig. 2.**
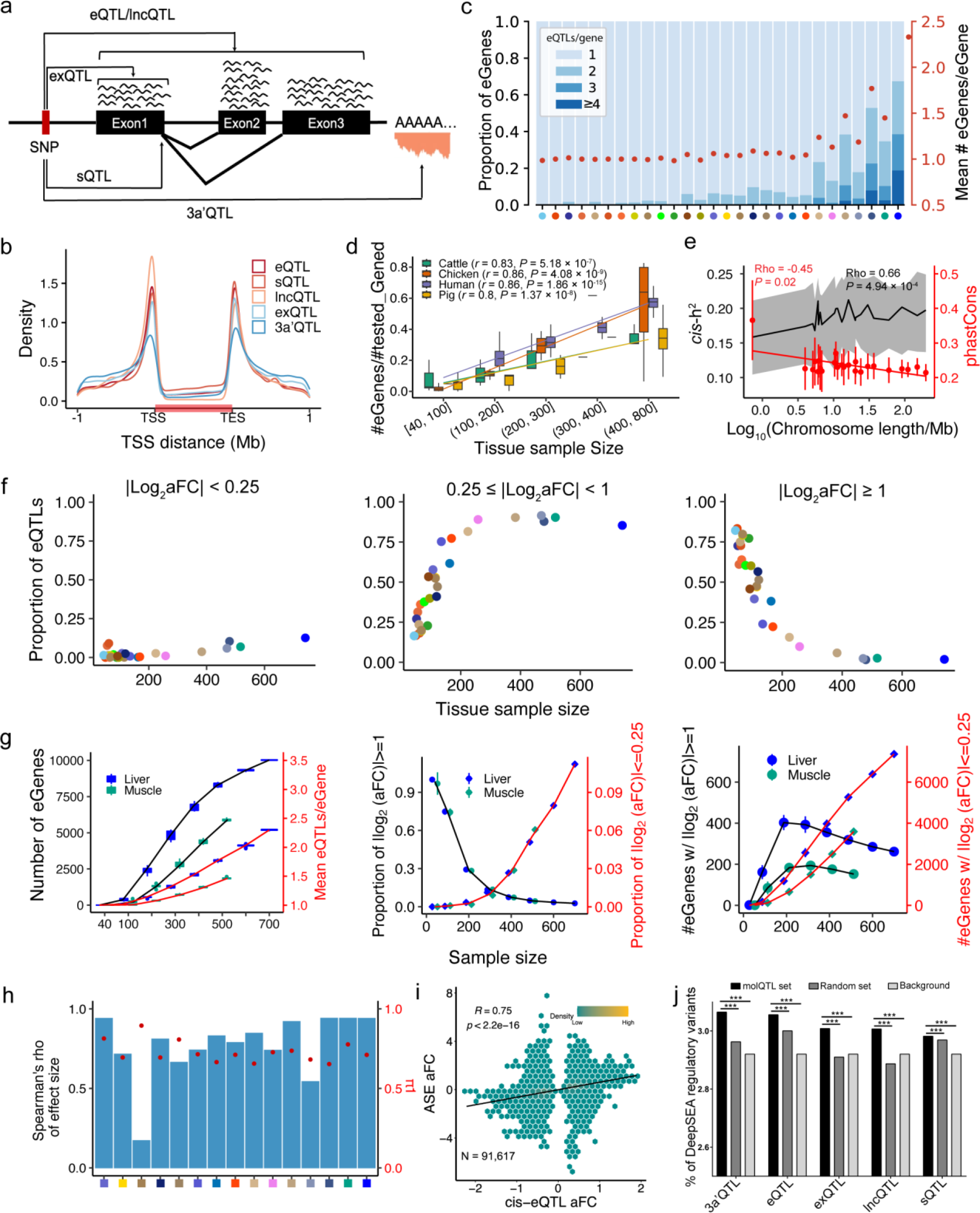
Molecular QTL (molQTL) mapping in 28 chicken tissues. (**a**) Illustration of the definition of five molecular phenotypes and the respective molQTL, including protein-coding gene expression (eQTL), lncRNA expression (lncQTL), exon expression (exQTL), splicing variation (sQTL), and 3’ untranslated region alternative polyadenylation (3’UTR APA, 3a’QTL). (**b**) Distribution of molQTL around gene body of eGenes, denoted by the horizontal red bar. TSS: Transcription Start Site, and TES: Transcription End Site. (**c**) Conditionally independent eQTL across all 28 tissues. Proportion of eGenes with different numbers of independent eQTL being detected (blue stacked bars; left *y*-axis), and mean number of independent eQTL per eGene (red dots; right *y*-axis). Tissues are sorted from smallest to largest regarding sample size. Tissue color legend can be found in Fig. 1c and Table S2. (**d**) Proportion of eGenes detected as a function of tissue sample size across species, including 28, 34, 24 and 49 tissues in chickens, pigs, cattle and human, respectively. The lines are fitted with a linear model implemented in the geom_smooth function of the ggplot2 package(*104*). Correlations and *P* values were computed with the Spearman method using the cor.test function in R v3.6.5(*105*). (**e**) *cis*-*h*^2^ (cis- heritability, left *y*-axis) and phastCons scores (right *y*-axis) of lead eQTL as a function of chromosome size (log10scaled). The top and bottom boundaries of the grey shade indicate the 25% and 75% of *cis*-*h*^2^ range, respectively, and the black line is the median of *cis*-*h*^2^ values. Red dots are average phastCons of lead eQTL, and red bars are their standard deviations. The correlations were computed with the Spearman method, and *P* values were computed *via* the asymptotic *t* approximation. (**f**) The proportion of eQTL detected (*y*-axis) with different effect sizes (from left to right panels) as a function of tissue sample size (*x*-axis). (**g**) Down-sampling analyses of eGene and eQTL. We carried out down-sampling analyses (10 replications at each sample size) in the liver and muscle, which have the largest sample size among all the 28 tissues. The left panel depicts the number of eGenes (left *y*-axis) and mean eQTL per eGene (right *y*- axis) detected at different sample size. The middle panel shows the proportion of detected eQTL of large (absolute log2aFC ≥ 1, left *y*-axis) and small effect size (absolute log2aFC ≤ 0.25, right *y*- axis). The right panel presents the number eGenes detected when the regulatory effect size of lead eQTL is large (absolute log2aFC ≥ 1, left *y*-axis) and small (absolute log2aFC ≤ 0.25, right *y*-axis). (**h**) Internal validation of eQTL. Bars in light blue indicate the Spearman correlation coefficient of eQTL effect size between validation and discovery groups (left *y*-axis), and red dots represent π1 statistic estimating the replication rate of eQTL between groups (right *y*-axis). The samples in each of the 15 tissues with over 100 individuals are evenly and randomly divided into two groups, i.e., discovery and validation groups. The tissue color legend (*x*-axis) can be found in Fig. 1c and **Table S2**. (**i**) Correlation between effect size of eQTL (*x*-axis, n=91,617) and those of same loci derived from allele-specific expression (ASE, *y*-axis) analysis in liver. (**j**) The proportion of regulatory variants predicted by DeepSEA (prediction score > 0.7) based on 310 functional profiles in chickens. molQTL_set: conditionally independent molQTL across tissues; Random_set: randomly selected variants with the same MAF as molQTL; Background: all tested 1.5 million variants.

The statistical power of molQTL mapping depends on the sample size of the tissue, similar to findings in other species(*32*, *47*, *48*) (**fig. 2d-g, figs. S13** and **14**). The down-sampling analysis in the liver and muscle further confirmed the relationship between sample size and eQTL discovery power (**fig. 2g**). Most eQTL with large effect (i.e., fold change of expression, aFC > 2) were detectable when sample size reached around 200 (**fig. 2f** and **g**), and eQTL with larger effect were more enriched around TSS (**fig. S14l**) and had lower minor allele frequency (MAF) (**fig. S14**). In general, the estimated effect size of eQTL was not correlated with their gene expression levels across tissues (**figs. S15 a** and **b**). Of note, chromosome size was significantly and positively correlated with eGene heritability (**fig. 2e**), eGene discovery (**fig. S15c**), and MAF of lead eVariants (**fig. S15d**). This was only observed in chickens and not in mammals (i.e., pig, cattle and human) (**figs. S15e-g**). Such influences of chromosome size on eQTL effects might be due to differences in evolutionary constraints between microchromosomes and macrochromosomes in chickens(*49*), which was further supported by the observation that phastCons scores of lead eVariants were also negatively correlated with chromosome size (**fig. 2e**).

To validate molQTL identified above, we first applied linear mixed model (LMM), by which we observed that the effect size and significance level of genetic variants estimated by the LMM were highly correlated with those estimated by the linear regression, implemented in tensorQTL (*50*) (**fig. S16**). We then randomly and evenly divided samples from 15 tissues with a sample size of over 100 into two subgroups, and then carried out eQTL mapping separately. A high replication rate, measured by π1(*51*), was observed between subgroups across tissues, ranging from 0.61 in the hypothalamus to 0.92 in the embryo (**fig. 2h**, **fig. S17**). The effect size of eQTL also exhibited a high Spearman’s correlation (an average of 0.77 across tissues) between subgroups (**fig. 2h**). Moreover, we observed that effect sizes derived from the eQTL mapping were positively and significantly correlated (an average of 0.52 across tissues) with those from the allele-specific analysis at the same loci (**fig. 2i, Table S8**). Finally, we trained a deep learning model of regulatory effects based on 310 functional epigenomic profiles in chicken *via* DeepSEA (*50*) (**fig. S15h**, **Table S7**), and observed that regulatory variants predicted by DeepSEA were more significantly enriched in eVariants than the expected (**fig. 2j**). Altogether, these results demonstrated the reliability of molQTL identified in this study.

### Limited sharing of regulatory mechanisms underlying five molQTL types

Out of all 27,203 tested genes, 16,097 (59.2%) had significant QTL for at least two molecular phenotypes (**fig. S18a**). The LD of lead variants of any two molQTL types from the same genes was low, ranging from 0.04 (exQTL *vs.* 3a’QTL) to 0.29 (exQTL *vs.* lncQTL) (**fig. 3a**). The colocalization analysis further confirmed the limited sharing of regulatory control among these molecular phenotypes (**fig. 3a**), indicative of their distinct genetic regulatory mechanisms. **Fig. 3b** takes *NLRC5* as an example, four molecular phenotypes of which were controlled by distinct genomic loci, and LD between the respective lead variants was lower than 0.07 (**fig. S18b**).

**Fig. 3.**
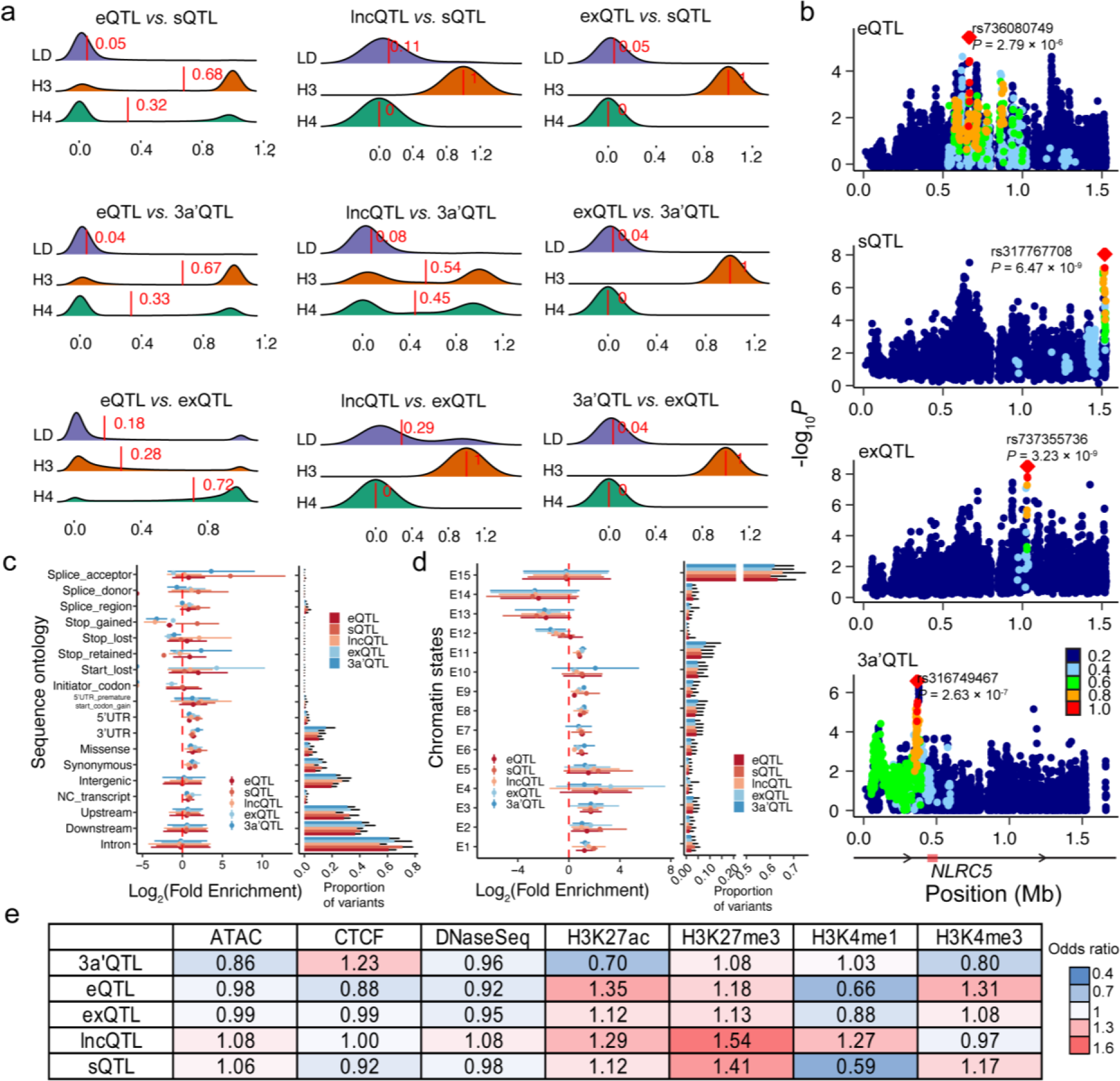
Colocalization and functional enrichment of molQTL. (**a**) Colocalization analyses of different types of molQTL of the same genes. The “LD” is the linkage disequilibrium (LD) of lead SNPs of two molecular phenotypes. “H3” and “H4” represent the probability of whether the association of two molecular phenotypes is due to two independent SNPs or one shared SNP, respectively. The vertical red lines indicate corresponding mean values. (**b**) Associations (i.e., - log10 transformed *P*) of genetic variants with four molecular phenotypes of *NLRC5*. The panels from top to bottom represent gene expression, alternative splicing, exon expression and 3’UTR APA, respectively. Color legend represents the degree of LD between the lead SNP and the others. The proportion and enrichment of five types of molQTL across sequence ontology (i.e., variant types annotated by SnpEff software(*106*)) (**c**) and 15 chromatin states (**d**). Fold enrichment is shown as mean (dot) ± standard deviation (log2 scaled, error bar) across 28 chicken tissues. The chromatin states were retrieved from Pan et al. (2023) (*30*). (**e**) The enrichment fold (odds ratio, OR) of molQTL in regulatory variants of seven epigenomic marks predicted by DeepSEA(*107*) (prediction score > 0.7). OR = (A/B)/(C/D), where C is the length of molQTL overlapped with annotated features (A), and B is the length of molQTL overlapped with the total genome length (D).

Among these molQTL, eQTL and exQTL exhibited a relatively high colocalization probability (average H4 = 0.72) (**fig. 3a**).

To elucidate molecular mechanisms of action behind these molQTL, we examined sequence ontology and multi-omics data in chickens, including 15 chromatin states predicted from 377 epigenetic data sets in 23 tissues(*30*), and 9,898 topologically associating domains (TADs) detected from high-throughput chromosome conformation capture (Hi-C) in three tissues (i.e., muscle, liver and testis)(*52*). As expected, conditionally independent molQTL were significantly enriched with various regulatory DNA sequences, including synonymous variants (1.67-fold in lncQTL to 3.03-fold in exQTL), 5’UTR variants (1.82-fold in eQTL to 3.64-fold in sQTL), 3’UTR variants (2.29-fold in sQTL to 3.77-fold in 3a’QTL), and non-coding (NC) transcripts (1.48-fold in 3a’QTL to 2.36-fold in exQTL). Of note, all five types of molQTL also exhibited a significant enrichment in missense variants (**fig. 3c**), indicating that a fraction of transcriptional regulatory variants may also alter protein amino acid residues(*53*). Compared to other molQTL, sQTL exhibited a higher enrichment with splicing variants (63.97-fold in splice acceptor, 3.97- fold in splice donor, and 3.89-fold in splice region), while 3a’QTL were more enriched with stop retained (5.06-fold) and 3’UTR variants (3.77-fold) (**fig. 3c**).

All five types of molQTL showed the highest enrichment in promoter-like states (E1-E5, an average of 3.64-fold), followed by enhancer-like states (E6-E10, an average of 1.98-fold) and ATAC islands (E11, an average of 1.87-fold). In contrast, they were significantly depleted from repressed regions (E12-E14) (**fig. 3d**). Compared to active enhancer (E6), super-enhancer (i.e. a cluster of enhancers in close genomic proximity, exhibiting exceptionally high levels of H3K4ac signals(*30*)), had a lower enrichment for all five molQTL, suggesting that they may be under a stronger purifying selection due to their essential roles in gene regulation and cell identity (**fig. S19a**). Among the five types of molQTL, 3a’QTL had the highest enrichment in enhancer-like states and ATAC islands (**fig. 3d**), supporting their high tissue-specificity. A total of 20% of eQTL, 26% of sQTL, 3.4% of lncQTL, 17.9% of exQTL, and 14.5% of 3a’QTL were supported by regulator-gene pairs that were predicted based on the correlation of signal density of regulators and gene expression (**fig. S19b, Table S9)**. By examining 3D looping of chromatin(*52*), we found 20-60% of molQTL-gene pairs located with the same TAD across tissues (**figs. S19 c** and **d**), with the highest enrichment observed at ∼400-600kp away from TSS of target genes after accounting for their distance (**fig. S19e**). As expected, 3a’QTL showed the highest enrichment at ∼600-1000kb downstream of their target genes (**fig. S19e**). Likewise, 41- 73% of eQTL-eGene pairs located in the same CTCF-loops that were identified from 22 chicken tissues (*30*) (**figs. S19f-h**). These results indicate that the long-distance eQTL exert effects possibly through disrupting TFBS in long-distance enhancers that interact with promoters *via* 3D looping of chromatin (**figs. S19c-h**). As shown in **fig. S19i,** eVariant *rs317368746* regulates expression of *TIMM17B* in the brain only, and it resides in a brain-specific enhancer and is located within the same TAD (346kb upstream) as the TSS of *TIMM17B*. Altogether, these results indicate that regulatory variants exert widespread effects on the transcriptome *via* multiple mechanisms such as changing transcript structure, function, stability, transcription/translation rate and chromatin conformation.

### Tissue- and breed-sharing of molQTL

All five types of molQTL were either tissue-specific or ubiquitous, among which 3a’QTL and eQTL exhibited the highest and lowest tissue-specificity, respectively (**fig. 4a** and **b, fig. S20a, fig. 21)**. This was also supported by the meta-tissue analysis (**fig. S22)**. In total, 10.6% of eQTL, 32.1% of sQTL, 27.4% of lncQTL, 25.8% of exQTL, and 29.6% of 3a’QTL were active in one tissue only. Of note, eQTL that were active in more tissues showed a higher enrichment around TSS (**fig. 4c**, **fig. S20b**), a smaller effect size (**fig. 4d**) and a higher MAF (**fig. S20c**). Tissue-shared eQTL (i.e., active in at least two tissues, LFSR < 0.05) also tended to be more enriched for promoter-like states, whereas tissue-specific eQTL were more enriched for enhancer-like states (**fig. S20d**). In general, tissues with similar biological functions (e.g., immune tissues) tended to be clustered together based on eQTL effect correlation (**fig. 4a, fig. S21**), which was similar for the remaining four types of molQTL (**fig. 4b, fig. S21)**. Unlike GTEx in mammals(*32*, *47*, *48*), blood formed the primary outgroup in chickens regarding eQTL and lncQTL, while, for the remaining three types of molQTL, brain and testis were first separated from the rest of the tissues. **Fig. S20e** demonstrates an eQTL (9_16035177_G_A) that significantly regulated the expression of *ALG3* only in the blood. The *ALG3* gene encodes alpha-1,3-mannosyltransferase with the function of inducing glycosylation of TGF-β receptor II(*54*), which might modulate blood pressure homeostasis(*55*) and affect hematopoiesis(*56*). In contrast to eQTL shared in other tissues, blood-specific eQTL had a lower MAF (**fig. S20f**) and a larger effect (**fig. S20g**). Moreover, genetic regulation of all five molecular phenotypes in the embryo was distinct from those in the primary tissues (**fig. 4a, fig. S21**), similar to that in pigs(*47*), indicating a distinct regulation of early development. In addition, we detected 59 eQTL with opposite directional effects on the same genes (n = 51) between tissues (**Table S10**). For instance, the T-allele of *rs315639985* increased the expression of *FBXO5* in the spleen but decreased its expression in the whole blood. The *FBXO5* gene encodes F-box protein 5, which is associated with systolic blood pressure in human(*57*) (**fig. S20h**). Another example was *rs313608694,* whose G-allele significantly upregulated the expression of *ELAC2* in the embryo but downregulated it in the spleen (**fig. S20h**). This gene encodes elaC ribonuclease Z 2, and the reduction of its expression could induce growth arrest by suppressing transforming growth factor-beta(*58*).

**Fig. 4.**
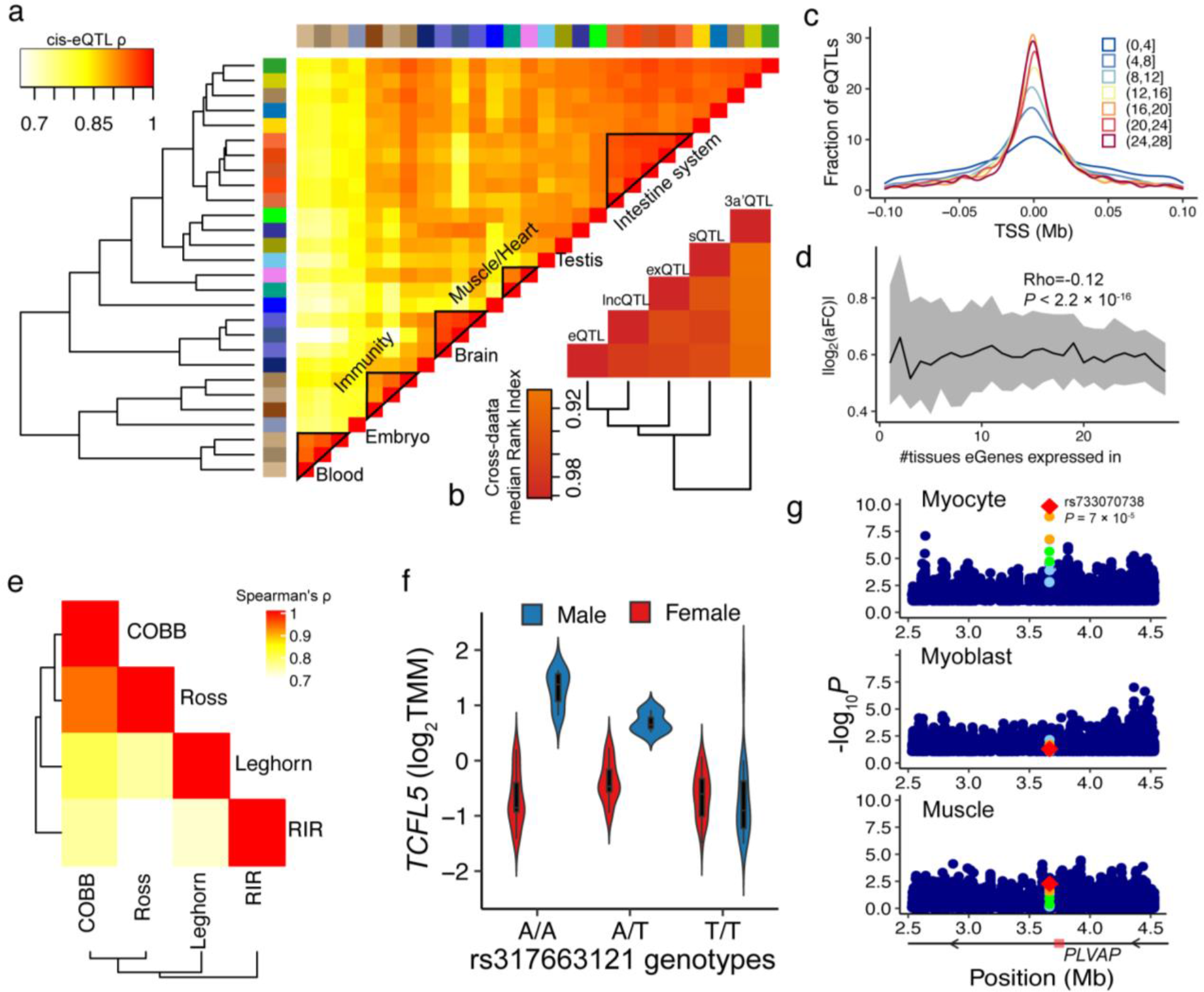
Tissue-sharing and context-dependent patterns of molQTL. (**a**) The heatmap of Spearman’s correlation of eQTL effect size between tissues. Tissues are clustered on the basis of dissimilarities (i.e. 1-d), where d is Euclidean distance calculated from the eQTL effect, with a complete linkage method(*108*). The color legend of tissues is the same as in Fig. 1c and **Table S2.** (**b**) Similarity of molQTL effect-based tissue clustering patterns. The pairwise Rand Index across five types of molQTL is used for measuring the similarity, ranging from 0 to 1, where 0 means that two tissue clustering patterns do not match at all, while 1 means that two clustering patterns match exactly. (**c**) Fraction of eQTL around transcription start site (TSS) according to number of tissues they are active in. (**d**) Absolute effect size (allelic fold change, aFC) of eQTL as a function of number of tissues where the eGene is expressed in. The black line is corresponding median estimates, and the grey shades indicate corresponding interquartile ranges. Correlation tests were carried out using *cor.test* function in R v3.6.3. (**e**) Heatmap depicting of eQTL effect sharing between breeds. This analysis was done by using Multivariate Adaptive Shrinkage (*109*), same as in panel **a**. (**f**) The expression of the *TCFL5* gene across three genotypes of *rs317663121* in males (AA, n=107; AT, n=106; TT, n=92) and female (AA, n=66; AT, n=56; TT, n=73). TMM is the Trimmed Means of M values, representing the normalized gene expression level. (**g**) The significance (-log10*P*) of interaction between *rs733070738* genotypes and myocyte enrichment (top panel) and myoblast enrichment (middle panel) on *PLVAP* expression in muscle. The bottom panel is the Manhattan plot for eQTL mapping of *PLVAP* in bulk muscle samples. Dot color means linkage disequilibrium (LD) between *rs733070738* and the rest.

We examined breed-sharing of eQTL in the brain, liver, muscle and spleen, as all of them had more than two breeds and each with a sample size > 40. As a result, the majority of eQTL (an average of 81%) could be replicated between breeds and the replication rate was associated with tissue sample size (**fig. S23a, Table S11**). Furthermore, the eQTL effect was substantially shared between breeds (**fig. 4e**). For instance, the T-allele of *rs314795649* significantly upregulated expression of *PRKCDBP* in the liver across all four breeds being tested, including Cobb (β = 0.57, *P* = 2.67 × 10^-6^), Leghorn (β = 0.33, *P* = 3.10 × 10^-6^), Rhode Island Red (β = 0.39, *P* = 3.06 × 10^-6^) and Ross (β = 0.37, *P* = 5.02 × 10^-10^) (**fig. S23b**). In addition, we detected 376 (Red Jungle Fowl vs. Ross) and 185 (Red Jungle Fowl vs. Leghorn) breed-interaction eQTL (bi- eQTL) in the brain, and with genes regulated by them were enriched in functionals related to brain development (**Table S12**).

### Context-dependence of molQTL

To explore the context-dependent nature of gene regulation, we systematically detected eQTL interacting with sex (sb-eQTL), transcription factor (TF-eQTL) and cell type (ci-eQTL). For sb-eQTL mapping, we only considered eight tissues, where each sex had data from over 30 individuals available. In total, 1,138 SNPs displayed sex-biased regulation of 962 eGenes (sb- eGene, FDR < 0.01), ranging from 3 in the small intestine to 954 in the liver (**URL**). Taking the liver as an example, we further performed the sb-eQTL mapping in a single breed, Rhode Island Red (nmale= 32; nfemale= 46), resulting in 48 sb-eQTL regulating 30 eGenes (**fig. 4f, figs. S23c-d, Table S13**). For instance, the significant association of *rs317663121* with *TCFL5* expression was only observed in male liver (**fig. 4f**). Moreover, 14% (164) of sb-eGenes overlapped with sex-biased expressed genes in all eight studied tissues. These sb-eGenes detected in the blood, hypothalamus, and liver were significantly enriched in biological processes related to amino acid metabolism, signaling transduction pathway, and fatty acid metabolism (**Table S14**). Through the examination of 956 chicken transcription factors retrieved from the AnimalTFDB 3.0(*59*), we detected an average of 1,941 TF-eQTL in 17 tissues, representing 503 TFs (**fig. S23f, URL**). **Fig. S23e** illustrates that effect of *rs313600592* on *ATP6V1A* expression was significantly associated with the expression of transcription factor *TCF25* in the muscle. For ci-eQTL mapping, we first annotated 13 cell types from single-cell RNA-Seq data in chicken heart and muscle (**Table S15**). Based on the cellular composition of bulk RNA-Seq samples of muscle and heart estimated by the *in silico* cell-type deconvolution (**fig. S24**), we identified an average of 105 ci-eGenes in the muscle, ranging from 11 with interactions in adipocytes to 214 with Schwann cells, and an average of 19 ci-eGenes in the heart, ranging from 6 interacting with fibroblasts to 36 with cardiomyocytes (**fig. S23g, Table S16**). For instance, *rs733070738* regulated expression of *PLVAP* by interacting with myocyte enrichment in the muscle (**fig. 4g**). These results highlight the dynamics of genetic regulatory effects across distinct biological contexts.

### Interpreting genetic regulation behind complex traits and adaptive evolution

To show the potential of molQTL in understanding complex traits in chickens, we systematically integrated molQTL with GWAS results of 108 complex traits, including growth and development (n = 43), carcass (n = 41), egg production (n = 20), feed efficiency (n = 3), and blood biochemical index (n = 1) (**Table S17**). Enrichment analysis revealed that GWAS loci of all the traits were significantly enriched in all five types of molQTL (**fig. S25a**). Among them, the highest enrichment was observed for 3a’QTL (1.87±0.33), followed by sQTL (1.83±0.28), eQTL (1.81±0.30), lncQTL (1.59±0.27) and finally exQTL (1.56±0.32) (**fig. S25a**).

Furthermore, we applied four complementary methods to prioritize causal variants and genes underlying each GWAS loci, including fastENLOC-based colocalization, summary-data-based MR (SMR), single-tissue transcriptome-wide association study (sTWAS), and multi-tissue TWAS (mTWAS). Out of all 1,176 significant GWAS loci, 1,059 (90%) could be explained by at least one molQTL across 28 tissues (**fig. 5a, figs. S26** and **S27**). Of 896 colocalized GWAS loci, 59.9% were not colocalized with the nearest genes of lead GWAS variants, indicative of the regulatory complexity of complex traits (**fig. 5b, fig. S28a**). The number of colocalization events of a trait was determined by the statistical power of both GWAS and molQTL mapping (**fig. S26b-c**). Of all 1,176 GWAS loci, 0.8%, 0.9%, 5.2% and 1.4% were explained uniquely by eQTL, sQTL, exQTL and lncQTL, respectively. This result indicates that each type of molecular phenotype only had a limited contribution to complex traits at distinct levels of gene regulation (**fig 5a**, **fig. S29**). Taking the body weight gain from week 6 to 8 (WG6.8) as an example, sTWAS linked GWAS loci to 43 unique genes (34 protein-coding and 9 lncRNA genes) across 21 tissues (**fig. 5c, fig. S28c, Table S18**). Of them, the expression of the *KPNA3* (karyopherin subunit alpha 3) exhibited the strongest association with WG6.8 in the retina, followed by pituitary and heart (**fig. 5c**). Consistently, it has been documented that the knockdown of the *KPNA3* would restore photoreceptor formation in *Drosophila*(*60*). The highest colocalization between WG6.8 GWAS loci and molQTL of *KPNA3* was observed for a retina eQTL (*rs314814283*, GRCP=0.78) and a pituitary sQTL (*rs13552958*, GRCP=0.54) (**Table S19**). The further SMR analysis pinpointed 10 potential causal mutations across tissues (**Table S20**), among which *rs739579746* was the most significant one (**fig. 5c, Table S20**). The SNPs *rs314814283* and *rs739579746* detected by eQTL mapping were in high LD (*r*^2^ = 0.88), while both showed low LD with *rs13552958* (r^2^ < 0.02) detected by sQTL mapping. These findings likely reflect the importance of photoreception for chicken growth and production performance (*61*, *62*), and the promising candidate gene in this region is the *KPNA3.* In addition, we detected 149 significant lncRNA-protein-coding-trait regulation events with SMR-multi analysis (**Table S21**). For instance, an eQTL of a lncRNA (*ENSGALG00000053557*), located on the opposite strand of the *IL20RA*, exhibited significant colocalizations with an eQTL of *IL20RA* in the muscle and GWAS loci of the total stomach weight on chromosome 3 (**fig. 5d**).

**Fig. 5.**
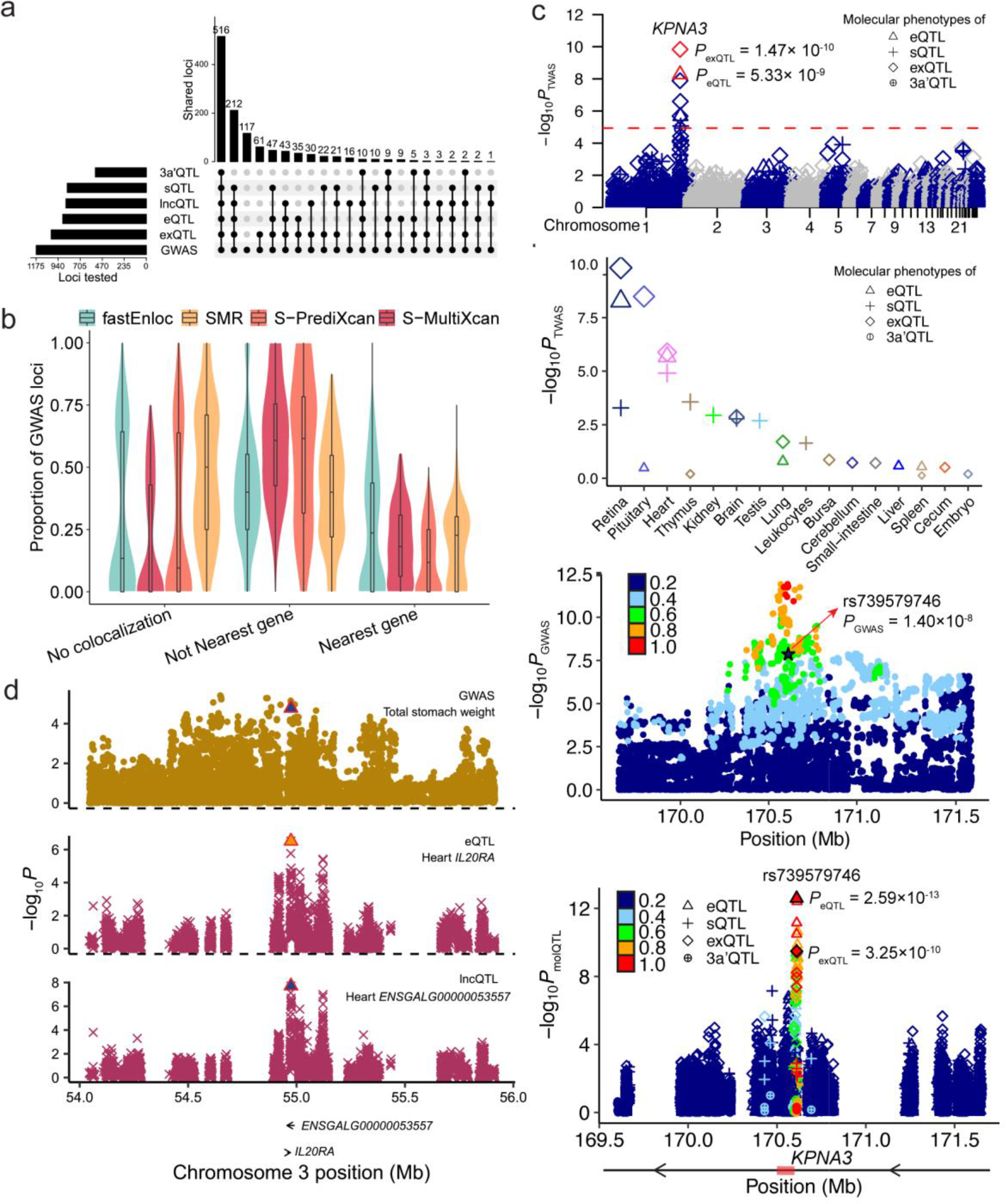
**Interpretation of GWAS loci with molQTL**. (**a**) UpsetR plot depicting the number of GWAS loci explained by five types of molQTL, which were detected by at least one of four complementary integrative methods, including fastENLOC-based colocalization, Summary- based Mendelian randomization (SMR), single tissue-based transcriptome-wide association study (sTWAS), and multi-tissue TWAS (mTWAS). (**b**) The proportion of three types of GWAS loci (n = 1,155) colocalizing with eQTL regarding the integration results using 4 methods, including fastENLOC-based colocalization, Summary-based Mendelian randomization (SMR), single tissue-based transcriptome-wide association study (sTWAS), and multi-tissue TWAS (mTWAS). No colocalization: GWAS loci that are not interpreted by any eGenes in 28 tissues. Not nearest gene: GWAS loci are interpreted by eGenes that are not nearest genes to GWAS lead SNPs. Nearest gene: GWAS loci are interpreted by eGenes that are the nearest ones to GWAS lead SNPs. Each dot represents one of 108 complex traits. (**c**) Interpretation of GWAS loci of weight gain from week 6 to 8 (WG6.8) with molQTL. The top panel depicts associations of genes with WG6.8 *via* sTWAS in retina. The second lower panel displays associations (-log10*P*) of different molecular phenotypes (gene expression, exon expression, alternative splicing and 3’UTR APA) of *KPNA3* with WG6.8 obtained by sTWAS across tissues. The following Manhattan plot exhibits GWAS associations of SNPs with WG6.8 on chromosome 1. The color indicated linkage disequilibrium (LD) of SNPs with the lead one (*rs15497848*, *P* = 1.22 ×10^-12^). The colocalized SNP (*rs739579746*, *P* = 1.4 ×10^-8^) is denoted as a black star. The bottom plot represents molQTL mapping results of *KPNA3* in retina. The color represents LD values of the colocalized SNP (*rs739579746*, black color) with the rest. (**d**) SMR-multi results of GWAS loci of total stomach weight and eQTL and lncQTL. The top panel depicts GWAS associations of SNPs (represented by dots) with total stomach weight. The middle panel exhibits SMR associations of GWAS loci with eQTL of *IL20RA* in heart, while the bottom panel exhibits SMR associations of GWAS loci with lncQTL of *ENSGALG00000053557*. The triangle shape shows the potential causal SNP (*rs314997637*) across the three biological layers, i.e., expression of *ENSGALG00000053557,* expression of *IL20RA* and total stomach weight.

To further explore context-specific genetic regulation of complex traits, we conducted colocalization analysis between GWAS loci and three types of context-interaction eQTL detected above (**fig. S25b**). Out of 1,155 GWAS loci, 22.9% (264), 48.7% (562) and 12.3% (142) were explained by sb-eQTL, TF-eQTL and ci-eQTL, respectively (**fig. S25b**). For instance, GWAS loci of total stomach weight and body weight at 8 weeks of age were significantly colocalized with sb-eQTL of *MFSD4A* and *TOX3* in the brain and spleen, respectively (**figs. S28d-e**). The *TOX3* gene encodes TOX high mobility group box family member 3, playing roles in sex determination and differentiation (*63*, *64*). Despite the limited discovery power of the context-interaction eQTL due to the small sample size, our analysis demonstrated that context-specific regulatory effects were nonnegligible in dissecting the regulatory mechanism of complex traits. Furthermore, we conducted an exploratory analysis to investigate whether domestication and breeding also tend to target on regulatory variants, though examining selection sweeps previously detected between broilers and layers previously (**fig. S25d, fig. S30**) (*14*, *65*). Within the brain, we separately detected eQTL in three chicken lines/breeds separately, including Red Jungle Fowl (n = 46), Ross (n=157) and Leghorn (n = 78). Genomic windows containing at least one eQTL (i.e., eQTL windows) in Ross and Leghorn were under stronger selection (i.e., larger selection values, LSBL) in broilers than expected, whereas those detected in Red Jungle Fowl were not (**fig. S25d, fig. S30**). Likewise, for selection sweeps in layers, eQTL windows in Leghorn were under stronger selection in layers than expected, but not for eQTL windows in Ross and Red Jungle Fowl (**fig. S25d, fig. S30**).

Altogether, the current ChickenGTEx can serve as a valuable resource for exploiting regulatory mechanisms underlying complex traits and adaptation in chickens.

### Comparing gene regulation and complex trait genetics between chickens and mammals

Based on gene orthology between chickens and three mammals (i.e., cattle, pigs and humans), we found the expression levels of the 1-1-1-1 orthologous genes were significantly higher than those of non-orthologous genes across tissues (**fig. S31)**. The proportion of orthologous genes expressed in chicken tissues was positively (Pearson’s *r* > 0.8, *P* < 0.004) correlated with that in mammalian tissues (**fig. S31c**). Based on gene expression profiles, 14,278 samples in the four species were clustered first according to their tissue types, indicating the global conservation of gene expression between chickens and mammals (**fig. 6a**). This was also supported by a high correlation of TAU values of genes, a measure of tissue-specificity of gene expression, between chickens and mammals (**fig. S31d**). The phylogenetic analysis of gene expression revealed different evolutionary rates of tissues across species, where testis and pituitary evolved fastest, while adipose and liver evolved slowest (**figs. S31e**). The effect sizes of lead eQTL of orthologous genes were significantly but weakly correlated between chickens and mammals, which were lower than those within mammals (**fig. 6b**). This was consistent for *cis*-*h*^2^ of orthologous genes (**fig. S32a**). As in pigs and cattle, the distance of lead eQTL to TSS was larger in chickens than that in humans (**fig. S32b** and **c**), which might be partially due to the larger LD of SNPs in farm animals’ genomes and lower SNP density in the pilot phase of FarmGTEx compared to human GTEx(*32*). We further divided chicken eGenes of each tissue into two groups: 1) chicken-specific eGenes, and 2) those shared with at least one mammalian species (conserved eGenes) (**see Methods**). In general, compared to chicken-specific eGenes, conserved eGenes showed a higher gene expression level, lower tissue-specificity, were more likely to be differentially expressed between species, have more promoters, and stronger tolerance to loss-of-function mutations (less evolutionarily constrained) (**fig 6c**).

**Fig. 6.**
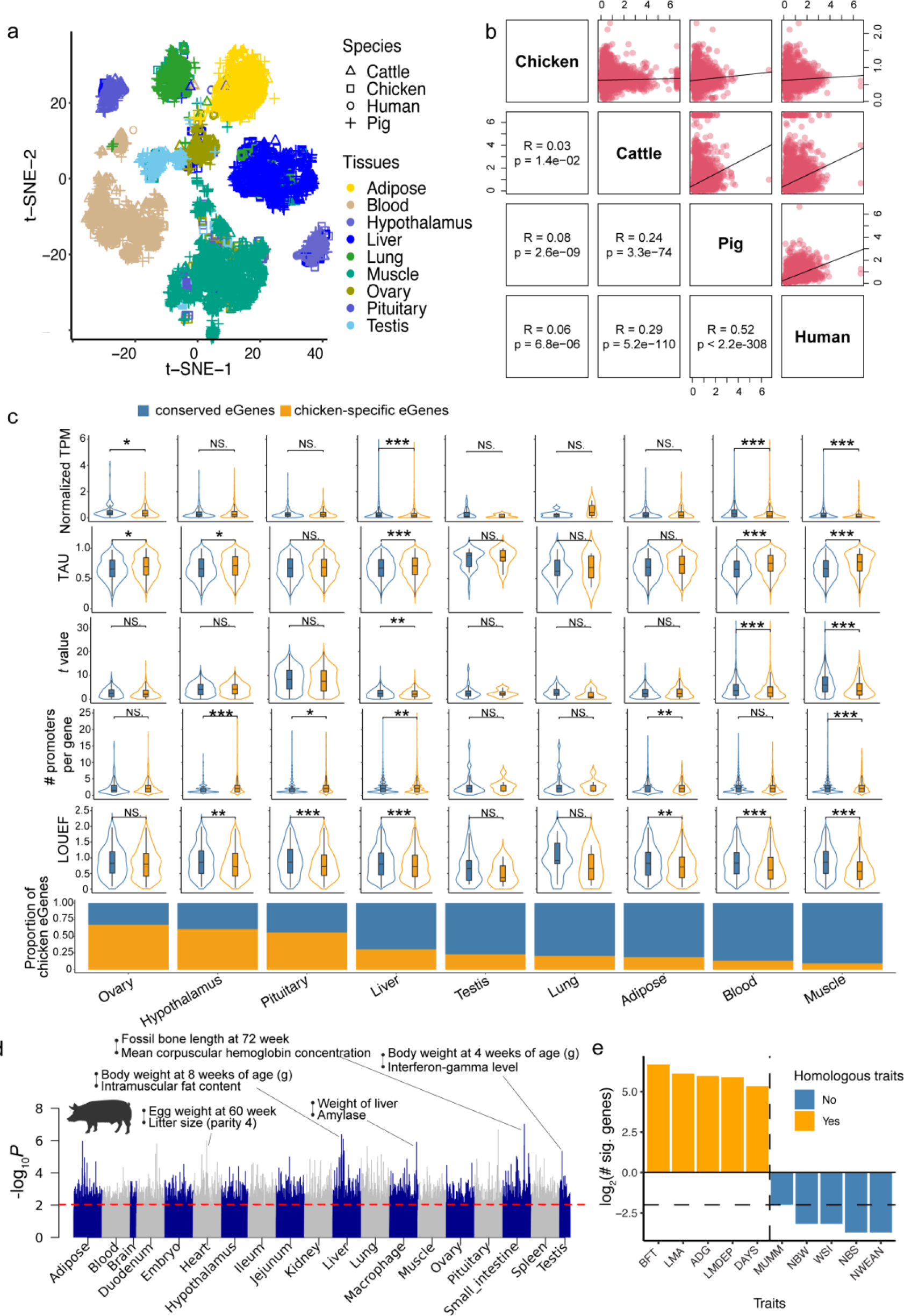
Comparative analyses of gene regulation and transcriptome-wide associations (TWAS) between chickens and mammals. (**a**) Visualization of variance in gene expression of 14,278 RNA-Seq samples across four species (i.e., Chicken, pig, cattle and human) *via* a *t*- distributed stochastic neighbor embedding (*t*-SNE). Gene expression (Transcripts per Million, TPM) of 10600 1-1-1-1 orthologous protein-coding genes are normalized between samples using Seurat software (v4.3.0) (*110*). (**b**) Pearson’s correlation of averaged effect size of lead eVariants of 5,513 orthologous eGenes between species. (**c**) Comparison of chicken-specific eGenes and conserved eGenes (i.e., eGenes that are shared with at least one mammalian species) across nine tissues. The bottom barplot depicts the proportion of chicken-specific eGenes and conserved eGenes in each tissue. The violin plots from top to bottom depict expression level, TAU (tissue-specificity of expression), *t*-value (measuring the degree of gene differential expression between species), number of promoters per gene, and loss-of-function intolerance (quantified by LOEUF), respectively. Statistical tests were done by employing two-sided Wilcox-test. \*\*\**P* ≤ 0.001; **0.001 < *P* ≤ 0.01; *0.01 < *P* ≤ 0.05; NS: not significant (*P* > 0.05). (**d**) Significance (at log10 transformed, *y*-axis) for TWAS-based correlations calculated using one-to-one orthologous gene effect between chicken and pig. The red dashed line depicts the threshold of significance (Permutation-based *P* value < 0.01, corresponding to nominal *P* value < 9.11 × 10^-3^). (**e**) The number of genes newly detected (FDR < 0.05) for body weight in chickens by using cross-species meta-TWAS analysis in muscle. The dashed horizontal line indicates 0 before the log transformation. Orange and blue bars represent homologous and nonhomologous traits in pigs for chicken body weight, respectively. BFT: backfat thickness, LMA: loin muscle area, ADG: average daily gain, LMDEP: loin muscle depth, DAYS: days, MUMM: number of mummified pigs, NBW: number of weak pigs, WSI: weaning to estrus interval, NBS: number of stillborn pigs, NWEAN: number of weaned piglets.

The FarmGTEx-based TWAS results provide new opportunities to systematically explore between-species similarity of complex trait genetics at the functional level of orthologous genes. We thus compared all the 3,024 sTWAS of 108 traits in chickens with 9,112, 1,032 and 6,480 sTWAS in three mammalian species, representing 268, 43 and 135 complex traits, respectively.

Within the matching tissues, we identified a total of 8,312 trait-pairs with significant correlations between chickens and three mammalian species (*P* < 9.11 × 10^-3^, permutation-based) (**fig. 6d**, **fig. S32d-f, Table S22**), despite the big differences in the TWAS power between species. Most of the significantly correlated traits between species recapitulated known biological and physiological knowledge. For instance, chicken body weight (BW) showed a high correlation with pig average daily gain (ADG) in the ileum (Pearson’s *r* = 0.69, *P* = 3.42 ×10^-5^, **fig. 32g**), cattle somatic cell scores (SCS) in the adipose (Pearson’s *r* = 0.38, *P* = 5.46 ×10^-5^, **fig. 32h**), and human type 2 diabetes (T2D) in the kidney (Pearson’s *r* = 0.57, *P* = 2.3 ×10^-5^, **fig. 32i**). This was in line with previous findings that larger BW fluctuation was related to an increased T2D risk in human(*66*), and also was positively associated with SCS in cattle (*67*). The expression of *ABCC13* (encoding ATP binding cassette subfamily C member 13) in the ileum was significantly associated with both chicken BW (*P* = 0.03) and pig ADG (*P* = 0.04), which encodes ATP binding cassette subfamily C member 13, which had potential associations with body weight/body mass index in humans (*68*). The expression of three genes, *PIGX*, *MRPL51* and *ABHD14B*, in the adipose were significantly associated with both chicken BW and cattle SCS. Of these, the ABHD14B protein is a lysine deacetylase with the capacity of catalyzing the deacetylation of lysine residues to yield acetyl-CoA, which could significantly alter glucose metabolism and could thus cause significant BW loss(*69*, *70*). The expression of *GABRB2* and *SOX4* in the kidney was significantly associated with both chicken BW and human T2D. The SOX4 is involved in pancreas development with roles in inhibiting insulin secretion and increasing diabetes risk(*71*, *72*). Moreover, taking chicken BW as an example, we carried out cross-species meta-TWAS analysis in the muscle, and found that homologous traits (e.g., ADG and back fat thickness) rather than non-homologous traits (e.g., number of stillborn and weaned pigs) in pigs could help detect more genes associated with BW in chickens **(fig. 6e)**. Similarly, human height and BMI increased the detection power of BW-associated genes in chickens *via* cross-species meta-TWAS analysis in the muscle (**fig. S32k**). These results highlighted that the FarmGTEx resource could facilitate the translation of genetic findings between species at the functional level of orthologous genes rather than the DNA sequence level.

## Discussions

### Summary and general impacts

Through the comprehensive analyses of the so-far largest collection of chicken RNA-Seq and WGS data, we have developed a catalogue of genetic variants with regulatory effects on five transcriptional phenotypes, representing both primary expression (including protein-coding, lncRNA and exon) and post-transcriptional modifications (alternative splicing and 3’UTR APA), across 28 chicken tissues, referred as the ChickenGTEx. We made the findings and resources of ChickenGTEx freely accessible to the entire community through http://chicken.farmgtex.org. This web portal provides an open-access chicken genotype imputation reference panel, which was built-up and maintained as part of this project. The current reference panel consists of approximately 3,000 WGS samples from around the globe, enabling researchers to impute genotypes derived from RNA-Seq, SNP array or low-coverage sequences to the whole-genome sequence level, which can be utilized further to prioritize potential causal variants underlying complex traits of interest through integrating with muti-layer ChickenGTEx resources. Besides, we offer highly-useful visualization tool, Integrative Genomics Viewer (IGV) (*73*) for exploring molecular phenotypes, enhancer-gene interactions, chromatin states, epigenetic modifications, and publicly available GWAS results. The web portal also includes single-cell RNA-Seq data that were collected and analyzed from six chicken tissues, enabling users to query the expression of their desired genes at both the cellular and bulk tissue level. Additionally, we provide batch data download and advanced search options for data resource generated in this study, and will continue updating the database to ensure its future accuracy and relevance. Overall, this first GTEx resource in avian species serves as a valuable resource for a global atlas of regulatory variants in chickens and informs vertebrate genome evolution at the functional level, benefiting future research in animal, plant, and human genetic and biomedicine research.

### MolQTL mapping and the underlying molecular mechanism

We have demonstrated that different molecular phenotypes of the same genes were likely to be controlled by distinct genomic loci through distinct regulatory mechanisms, indicating the importance of integrating omics data corresponding to multi-layer biologically-important molecular phenotypes (e.g., epigenetic mark activity and microRNA expression(*74*)) in future studies. This is consistent with findings in humans that most of the sQTL and 3a’QTL were distinct from eQTL (*75*, *76*). The comparative analysis of regulatory variants reveals several specificities of gene regulation in chickens compared to mammals. For instance, the chicken genome exhibits a chromosome-size dependence in genetic control of gene expression, in contrast to mammals. Avian genomes often have chromosomes of highly variable sizes, with chicken chromosomes ranging from a minimum of 3.4 Mb to a maximum of 200 Mb (*12*). Chicken microchromosomes exhibit a higher gene density, higher GC content and DNA methylation levels(*12*, *77*), and are under stronger evolutionary constraints (*49*), that easily distinguish them from the mammal ‘like’ macrochromosomes. These distinct genetic and epigenetic features might lead to differences in the genetic regulation of gene expressions across chromosomes in chickens. In addition, we observed a high sharing of eQTL effect across tissues in chickens, while interestingly the blood showed the highest dissimilarity against other tissue types. This observation is in contrast to that of mammals, where the testis showed the highest dissimilarity (*32*, *47*, *48*) that is perhaps a result of nucleated red blood cells in avian blood (*78*, *79*). Moreover, we uncovered a set of genetic variants with regulatory effects interacting with biological contexts, e.g., sex, transcription factor expression, genetic background, and cell type compositions. This context-dependent molQTL explained 10-50% of GWAS loci, revealing the need to consider cell types/states under different developmental stages, nutrition, and physiology status in the future molQTL mapping experiments. By taking account of a wide range of environmental/biological contexts, we can effectively tackle the challenge of “missing regulation” (*80*, *81*). As demonstrated in human studies(*82*, *83*), harmonizing data from diverse chicken breeds/lines increased the detection power of molQTL *via* increasing sample size, facilitating the fine-mapping of causal variants *via* reducing LD of SNPs, as well as allowing breed-specific molQTL mapping (*84*). At the current pilot phase, eQTL with *trans*-regulatory effect (> 1Mb to the TSS of genes) is not considered due to the limited sample size. Discovering *trans-*eQTL, which often has a small effect size, requires hundreds of thousands of samples (*82*, *85*), and will be considered in the future when the sample size of transcriptome data is sufficient.

### Potential applications of ChickenGTEx

This multi-tissue gene regulation resource opens the door to decipher the biological mechanism of complex traits, domestication and polygenic adaptation in chickens in-depth. It enables nearly 90% of GWAS loci being tested in this study to be explained by at least of one type of molQTL, a higher proportion than that in humans (78%) (*32*) or in pigs (80%) (*47*). This finding demonstrates the importance of molQTL mapping in functionally dissecting agriculturally important traits in farm animals, with a high potential for accelerating and improving the current animal breeding program and enabling the future precision selection and breeding (*26*, *86*). The focus of cross-species comparison studies in the past decades was mainly on the DNA sequence level due to the lack of relevant functional data, and the recent Zoonomia project investigated the DNA sequence evolution of regulatory elements while based on *in silico* prediction across species (*87–89*). The ChickenGTEx offers new means to explore the evolutionary impacts of gene regulation on complex traits across species and translate genetic findings between species at the functional level of orthologous genes rather than the DNA sequence level. Our exploratory comparative analysis of large-scale TWAS between chickens and mammals illustrates how to “borrow” information between species for gene mapping (*90*, *91*). We found that cross-species meta-TWAS aided in the identification of more functional genes for homologous traits. We believe that the ChickenGTEx resource will not only contribute significantly to elucidating the molecular architecture underlying phenotypic variation in chickens, but also to developing chicken models for studying human complex traits (e.g., disease and behavior (*3–7*)).

### Limitations and outlooks

The current ChickenGTEx provides the most expansive source of regulatory variants in the chicken genome. Some limitations and challenges remain in the genotype and molecular phenotype assessments. New chicken assemblies with more complete representation are becoming available with fewer computational limitations that we experienced using the GRCg6a reference genome (Ensembl version 102) (*12*, *92–94*). Future studies will consider long-read sequences to better resolve splice-variants (*95–97*), and pangenome references to annotate complex structural variants (*98*), mobile element variation (*99*), and short tandem repeats (*92*, *100*). In addition, it would be of great interest to investigate the functional impacts of rare and somatic variants on molecular phenotypes, where multi-tissue samples are collected from the same individuals with deep WGS data available. Beyond the bulk transcriptome, other molecular features could be included, e.g., DNA methylation variation, protein abundance, metabolite profiles, and the composition of the microbiome. For future single-cell genetics in chickens, a comprehensive chicken single-cell atlas will be the first step and is urgently required to explore the cell-type/state-specific gene regulation *via in silico* cell type deconvolution of large bulk tissue samples(*101*). In addition, conducting experimental follow-ups *via* methods, e.g., massively parallel CRISPR-based screens (*102*), is crucial to functionally validate and characterize regulatory effects of genetic variants and to identify functional genes of complex traits on a large-scale. In summary, the current and future versions of the ChickenGTEx project promises to establish a reference panel for studying the functional impacts of genetic variants in their native genomic and cellular contexts in distinct biological contexts, including molQTL mapping, molecular phenotype prediction for individuals with genotypes (including extinct species with ancient DNA available) and the evolution of regulatory variants. The fully developed ChickenGTEx will contribute substantially to research in complex-trait genetics, animal breeding, functional biology, and vertebrate genome evolution at the functional level.

## Acknowledgments

We thank all the researchers who have contributed to the publicly available data used in this research.

## Author contribution statement

L.Fang, H. Zhou, D.G., X.H., and N.Y. conceived and designed the project. D.G., Y.Y., B.Z. and Z.P. performed bioinformatic analyses of RNA-Seq data analysis. D.G., F.L., S.D., Y.G. and H.Y. conducted whole-genome sequence data analysis. D.Z., performed the deep learning analysis. D.G. performed multi-omics and single-cell RNA-Seq data analysis. D.G. conducted molQTL mapping. X.Z., C.Z. D.G. performed GWAS integrative analysis. Z.B. and D.G. led the comparison of GTEx between chickens and mammals. L.F., H.Zhou, D.G., X.Z., Q.L., C.Z., Y.H., Y.W., C.S., J.T., F.D., S.L., Y.W., M.W., M.P., D.R., M.C., J.S., K.W., A.J.B., W.W., L.Frantz, G.L., M.S.L., G.S., S.S., D.S., S.J.L., X.Z., B.L., H.Zhang, and H.C. contributed to the critical interpretation of analytical results before and during manuscript preparation. Y.H., D.Z., R.W., T.X., and Zhang Zhang built the ChickenGTEx web portal. H. Zhou, L.Fang, N.Y., X.H., G.E.L., Zhe Zhang, S.S., D.S., X.Z., Q.N., Z.L., W.L., H.Q., W. S. and C.L. contributed to the data and computational resources. D.G., Z.B., X.Z., C.Z., Y.W., Y.H. and L.Fang drafted the manuscript. All authors read, edited, and approved the final manuscript.

## Data and materials availability

All raw data analyzed in this study are publicly available for download without restrictions from SRA (https://www.ncbi.nlm.nih.gov/sra/) and NGDC BioProject (https://bigd.big.ac.cn/bioproject/) databases. Details of RNA-Seq, WGS, ChIP-Seq peaks and single-cell RNA-Seq can be found in Table S1, S6, S7 and S15, respectively. All processed data and the full summary statistics of molQTL mapping and genotype imputation reference panel are available at http://chicken.farmgtex.org. All the computational scripts and codes for RNA-Seq, WGS, single-cell RNA-Seq and Hi-C datasets analyses, as well as the respective quality control, molecular phenotype normalization, genotype imputation, molQTL mapping, functional enrichment, colocalization, SMR and TWAS are available at the FarmGTEx GitHub website (https://github.com/FarmOmics/ChickenGTEx_pilot_phase).

## Methods and Materials

### RNA-Seq data analyses and molecular phenotype definition

We downloaded 8,338 RNA-Seq data sets from the Sequence Read Archive (SRA, https://www.ncbi.nlm.nih.gov/sra)and 140 public data sets from the Genome Sequence Archive (GSA, https://ngdc.cncb.ac.cn/gsa/). We also included 155 newly-generated RNA-Seq data sets. The metadata relating to all the RNA-Seq samples is summarized in **Table S1**. For quality control, we removed adaptors and trimmed low-quality reads using Trim Galore (v0.6.6, https://github.com/FelixKrueger/TrimGalore) with options of “--gzip --trim-n --length 30 -- clip_R1 3 --clip_R2 3 --three_prime_clip_R1 3 --three_prime_clip_R2 3”. We aligned the clean reads to the GRCg6a reference genome (Ensembl version 102) using STAR (v2.7.7a)(111) with parameters of “--quantMode GeneCounts --chimSegmentMin 10 --chimOutType Junctions -- chimOutJunctionFormat 1 --outFilterMismatchNmax 3”. For downstream analyses, only 7,015 samples with uniquely mapping rates ≥ 60 % and a number of clean reads > 500,000 after removing potentially mislabeled samples were kept. For each of these samples, we then obtained raw read counts and normalized expression (i.e., Transcripts Per Million, TPM) of 16,779 PCGs annotated in the Ensembl v102 and 22,792 lncRNA genes annotated by FR-AgENCODE (http://www.fragencode.org/)(112), using featureCounts (v2.0.1)(*113*) and StringTie (v2.1.5)(*114*), respectively. Using the same software(*113*), we counted the total number of reads as a function of annotated exons, which were further transformed into TPM using TBtools(*115*). We performed the tree clustering of all the RNA-Seq samples using the GGTREE package(*116*). The distance between samples was measured by 1-*r*, where *r* was Pearson’s correlation coefficient based on the log2(TPM+0.25) of 5,000 genes with the highest variability. We also visualized these samples using the *t*-distributed stochastic neighbor embedding (*t*-SNE) approach implemented in the Rtsne package(*117*).

We quantified alternative splicing variation from RNA-Seq data using the LeafCutter package(*118*), which took into account spliced reads so that both novel and known alternative splicing events could be identified and quantified(*118*). Briefly, based on the STAR alignments mentioned above, we extracted junctions and defined intron clusters across samples using the script “bam2junc.sh” and leafcutter_cluster.py”, respectively, as provided by the LeafCutter package(*118*). For intron clustering, we required at least 30 split reads supporting each cluster and at least 0.1% of reads supporting a junction in a cluster, as well as allowing intron length of up to 500kb. The generated matrix of per individual counts was normalized and used for clustering samples based on 1-*r*, where *r* is the Pearson’s correlation coefficient between samples. To link intron clusters to genes, we mapped their coordinates to the gene model provided by the FR-AgENCODE database (*112*) using the script “map_clusters_to_genes.R” (https://github.com/broadinstitute/gtex-pipeline). Afterward, we filtered out introns if no reads were detected in >50% of samples or the number of counts was less than max(10, 0.1n) where n is the sample size. In addition, we discarded introns with low variability across samples: ∑i(|zi| < 0.25) ≥ n-3 and ∑i(|zi| > 6) ≤ 3, where zi is the z-score of the ^i^th cluster read fraction across individuals. The filtered counts were further normalized between samples using the script “prepare_phenotype_table.py” in the LeafCutter package(*118*). The generated normalized splicing counts were stored in BED formatted file for subsequent sQTL mapping.

For the quantification of 3’UTR APA, we utilized the DaPars (v2)(*119*). We first extracted distal polyadenylation sites based on the Ensembl annotation (v102) using the script “DaPars_Extract_Anno.py”. Then, we computed the genome coverage of STAR alignments mentioned above using the *genomecov* function in the BEDTools (v2.30.0)(*120*). The generated wiggle alignment files were then used for quantifying APA usage, resulting in the percentage of distal poly(A) site usage index (PDUI) value for each gene in each sample. We rescaled the PDUI values across samples to the mean of zero and variance of one in each tissue for 3a’QTL mapping.

### Single-cell RNA-Seq data analyses

We retrieved single-cell RNA-Seq data from the chicken heart (n = 7) (*121*) and muscle (n = 2) (*122*) from the public database. Raw sequencing data was processed by using the “count” function after preparing the genome annotation .gtf file (Ensembl v102) with the *mkgtf* tool of the Cell Ranger pipeline(*123*). The Seurat R package (v4.0.5)(*124*) was used for subsequent cell-type identification. We first created the Seurat object based on the raw read count of each sample in a tissue using the *CreateSeuratObject* function. In this step, we filtered out cells with unique gene counts < 200 and with mitochondrial counts > 20% of the total counts. We then normalized raw counts of gene expression using the *LogNormalize* algorithm and further identified highly variable genes (HVG) using the *FindVariableFeatures* algorithm with default parameters. The HVG count matrices of all samples for a given tissue were integrated and combined to form a single *Seurat* object using the *FindIntegrationAnchors* and *IntegrateData* functions. We scaled the integrated dataset using the *ScaleData* function, which was further used to run principal components analysis (PCA) with the *RunPCA* function. The top 15 PCs, where the percentage of variance explained tended to be constant based on the elbow plot by the *JackStraw* function, were selected for running Uniform Manifold Approximation and Projection (UMAP) analysis for cell clustering using the *RunUMAP* function. The nearest neighbors between cells were constructed using the *FindNeighbors* function and cell clusters were thus determined using the *FindClusters* function at a resolution of 0.05. We manually assigned cell names based on original publications (*121*, *122*) and the PanglaoDB database(*125*). Finally, cell clusters were visualized using the UMAP algorithm with the *DimPlot* function. To further deconvolute bulk RNA-Seq data using single-cell RNA-Seq data, we first created a signature matrix using the CIBERSORTx tool (*126*) with default parameters. Using the “Impute Cell Fractions” from the same tool, we imputed cell fractions with the custom mode and 1000 permutations after uploading the gene expression matrix for the bulk RNA-Seq data.

### Tissue-specificity of gene expression

We employed tspex(*127*), a tissue-specificity calculator, to compute 12 tissue-specificity metrics, including 8 general scoring metrics (i.e. Couts, Tau, Gini coefficient, Simpson index, Shannon entropy specificity, ROKU specificity, Specificity measure dispersion, and Jensen-Shannon specificity dispersion) and 4 individualized scoring metrics (i.e. Tissue-specificity index, Z-score, Specificity measure, and Jensen-Shannon specificity). To identify tissue-specifically expressed genes in a tissue, we applied another *t*-statistic approach as described previously(*128*). Briefly, for a given tissue, we carried out differential gene expression analysis between the target tissue and the rest but excluded those from the same biological system using the limma package(*129*). Subsequently, tissue-specific genes were identified when FDR corrected *P*- value(*130*) > 0.05 and fold change > 2. Functional enrichment analysis of tissue-specific genes with Biological Process (BP) terms in the Gene Ontology (GO) database was performed using the clusterProfiler package(*103*).

### Sex-biased gene expression

To identify genes with sex-biased expression, we employed DESeq2 software(*131*) to carry out differential expression analysis between male and female samples in 18 tissues, where sample size of each sex was > 10. We fitted a generalized linear model for the differential expression analysis while correcting for factors, including BioProject, year when RNA-Seq data was generated, age, breed, sequencing platform, library layout and selection method. After multiple testing correction by the FDR approach(*130*), the set of differentially expressed genes was identified when FDR corrected *P*-value < 0.01.

### Reference-guided transcript assembly

Based on STAR alignment files, we assembled transcripts with the guidance of the Ensembl annotation (GRCg6a v102) using the StringTie2 software tool(*114*). To increase the computational efficiency, transcript assembly was run by tissue. Then, the generated assembly files from all tissues were merged by using the “merge” function of the StringTie2 software(*114*). After quantifying the expression of assembled transcripts, we only retained single-exon transcripts with TPM >1 in at least half of samples in a tissue, and multi-exon transcripts with TPM >0.1 in at least half of samples in a tissue. Moreover, we compared our prediction to Ensembl and NCBI annotations using GffCompare (version 0.11), and classified them into 14 classes as described previously(*95*, *132*). The coding potential of predicted transcripts was predicted by using CPP2 software(*133*), with lncRNA loci predicted using FEElnc(*134*).

### Construction of gene co-expression networks

To build gene co-expression networks in each tissue, we employed five complementary methods with default parameters, including WGCNA (v1.69)(*135*), ICA (v1.0.2)(*136*), PEER (v1.3)(*137*), MEGENA (v1.3.7)(*138*), and CEMiTool (v1.8.3)(*139*). The input gene expression values were adjusted for hidden confounding factors by regressing out 10 PEER factors and 5 genotypic PCs (see “**Preparation for molQTL mapping**” section). Functional enrichment analysis of gene co-expression modules was conducted by using clusterProfiler (v4.0)(*103*), and the following visualization was done using Gephi (v0.9.2)(*140*).

### SNP calling from RNA-Seq samples

To call SNPs from RNA-Seq samples, we marked PCR duplicates in STAR alignment files and split reads that contained Ns in their cigar string using *MarkDuplicates* and *SplitNCigarReads* modules of the Genome Analysis Toolkit (GATK, v 4.1.9.0)(*45*), respectively. Using the Ensembl dbSNP database (v102), we recalibrated base quality scores using GATK *BaseRecalibrator* and *ApplyBQSR* modules. By following the best practice of germline variant calling from RNA-Seq data, we detected small variants from the recalibrated alignments files, which generated individual Genomic Variant Call Format (GVCF) files using the *HaplotypeCaller* function of the GATK tool(*45*). Then, we carried out joint-calling of all GVCF samples using the *GenotypeGVCFs* module from the GATK tool(*45*). For selecting high-quality SNPs, we carried out a hard-filtering with criteria of “FS > 30.0 & QD < 2.0”, resulting in a total set of 12,191,306 SNPs.

### Construction of the multi-breed genotype imputation panel and genotype imputation

We retrieved 1,693 public WGS data sets from SRA (n=1,213) and GSA (n = 480) databases along with 1,176 additional newly generated WGS samples, resulting in a total set of 2,869 WGS samples (**Table S6**). All raw sequence reads passed a uniform computational pipeline, including adaptor removal, read alignment, and SNP calling. Briefly, we trimmed read adaptors and low-quality reads using the Trimmomatic v0.39 software(*141*). The obtained clean reads were further aligned against the Ensembl GRCg6a chicken reference genome (v102) using the MEM algorithm of the Burrows-Wheeler Aligner (BWA, v0.7.17)(*142*). The alignment files in Binary Alignment Map (BAM) format were sorted using SAMtools (v1.9)(*143*), and were further passed for the removal of PCR duplicates using GATK (v4.1.9.0)(*45*). The obtained BAM files were then used for variant discovery to generate individual GVCF files using the *HaplotypeCaller* function of the GATK tool(*45*). The joint-calling of all 2,869 GVCF samples was further done using the *GenotypeGVCFs* module from the GATK tool(*45*). For selecting high-quality SNPs, we carried out a hard-filtering with criteria of “QD < 2.0, MQ < 40.0, FS > 60.0, SOR > 3.0, MQRankSum < -12.5, and ReadPosRankSum < -8.0”, resulting in a total set of 117,900,812 clean SNPs. To create the genotype imputation reference panel, we first filtered out multi-allelic and sex chromosomal SNPs, as well as those with MAF < 0.01 and missing rate > 0.9 using BCFtools v1.10.2(*144*), and then imputed missing genotypes using the Beagle 5.1 program(*145*). This yielded the final reference panel consisting of 2,869 samples and 10,520,420 SNP genotypes. To better impute SNPs called from RNA-Seq samples, we discarded SNPs called from RNA-Seq samples with MAF < 0.05 using BCFtools v1.10.2(*144*) and further evaluated the effect of missing rates decreasing from 0.9 gradually to 0.6 on imputation accuracy. This evaluation revealed that the missing rate of 0.6 could reach >95% of imputation accuracy, yielding a set of 1.5 million SNPs for subsequent analysis. The genotype imputation was performed using the Beagle 5.1 program(*145*).

### Preparation for molQTL mapping

*Sample deduplication.* After assigning RNA-Seq samples into 28 tissue types, we calculated the identity-by-state (IBS) distance between samples within each tissue based on the imputed SNP genotypes using PLINK v1.9(*146*). The formula of IBS calculation is as follows:

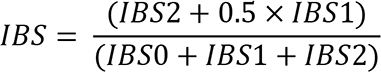

where IBS0, IBS1 and IBS2 are the number of non-missing variants when IBS = 0, IBS = 1, and IBS = 2, respectively. If the IBS distance of a pair of samples is higher than 0.9, they were deemed as duplicated so that the samples with the higher sequencing depth were kept. The deduplication process was run until all IBS values per pair of samples were less than 0.9. Finally, a total of 28 tissues with sample sizes ranging from 44 (testis) to 741 (liver) were kept for subsequent molQTL mapping.

*Principal component analysis*. Within each of those 28 tissues, we first LD-pruned imputed genotypes with the option of “--indep-pairwise 200 100 0.1” using PLINK v1.9(*146*). Principal component analysis (PCA) of samples was then carried out, based on the LD-pruned genotypes using the smartpca tool of the EIGENSOFT v8.0.0 package(*147*). The top 5 principal components (PCs) were selected as covariates for heritability estimation and molQTL mapping.

*Estimating PEER confounder factors*. To correct for confounders and other unwanted technical or biological variations in RNA-Seq samples, we estimated the Probabilistic Estimation of Expression Residuals (PEER) in each of the tissues using the PEER software package(*137*). The top 10 PEER factors showing highly relative contributions (i.e., factor weight variance) to gene expression variation were selected for subsequent heritability estimation and molQTL mapping.

*Phenotype preparation.* For protein-coding genes, lncRNAs and exons, we filtered out features with TPM < 0.1 and raw read counts < 6 in > 20% of samples within a tissue. Raw read counts were normalized using the Trimmed Mean of M-value (TMM) algorithm of the edgeR package(*148*). The generated TMM matrix was then further normalized with an inverse normal transformation for subsequent molQTL mapping. For splicing and APA, the preparation of molecular phenotypes was described in the “**RNA-Seq data analysis and molecular phenotype definition**” section.

### Estimating *cis*-heritability of gene expression

We leveraged the GCTA program v1.93.2(*149*) to estimate *cis*-heritability (*cis*-h^2^) of molecular phenotypes by fitting a mixed linear model:

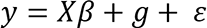

where y is a vector of phenotypic values (i.e., gene expressions) of all samples, *β* is a vector of corresponding coefficients of quantitative covariates X of all samples, which included 5 genotypic PCs and 10 PEER factors, g is a vector of the genetic values of SNPs around ±1Mb of the transcription start sites (TSS) of a gene, and ɛ is a vector of residuals. The genetic value *g* followed a normal distribution with mean of 0 and variance of ***A**σ*^2^_*g*_, where ***A*** was the genetic relationship matrix (GRM) between individuals(*150*). Thus, we can estimate *σ*^2^_*g*_, i.e., the variance explained by SNP genotypes (i.e., *cis*-h^2^), using the restricted maximum likelihood (REML) approach(*150*, *151*) implemented in GCTA software(*149*). The *cis*-h^2^ was finally defined when the significance level was lower than 5% based on the likelihood ratio test(*149*, *150*).

### Molecular QTL mapping

In this study, we only intended to map *cis*-molQTL of each feature, i.e., SNPs distributed around 1 Mb upstream and downstream of the TSS of the gene, using tensorQTL v1.0.4(*152*). This utilized graphics processing units (GPUs) with the scalability to increase runtime and reduce the time cost. Initializing with the option of “--mode cis_nominal” of the tensorQTL v1.0.4(*152*), we calculated all nominal associations of all variant-molecular phenotype pairs. The permutation mode was further used for computing empirical *P*-values for a molecular phenotype using the option of “--mode cis” of the tensorQTL v1.0.4. After carrying out a multiple testing correction based on empirical beta-approximated *P*-values(*153*) using the false discovery rate (FDR) approach(*130*), we defined eGenes, i.e., genes that were significantly regulated by at least one variant (FDR < 0.05). For an eGene, the empirical *P-*value that was closest to an FDR of 0.05 was defined as the genome-wide empirical *P*-value threshold (pt), which was used for defining the gene-level significance threshold using qbeta(pt, beta_shape1, beta_shape2) in R (v3.6.3)(*105*), where beta_shape1 and beta_shape2 were computed by tensorQTL v1.0.4(*152*).

The significant molQTL were tested SNPs whose nominal *P-*values were lower than the gene-level significance threshold.

### Fine-mapping analysis of molQTL

We employed four strategies for fine-mapping independent variants underlying each molQTL. Firstly, we utilized the stepwise regression procedure for mapping conditionally independent molQTL, as used in other GTEx studies (*32*, *47*, *48*). This analysis was done by using the tensorQTL v1.0.4 with “--mode cis_independent” option(*152*). The conditionally independent molQTL mapping was based on the nominal associations mentioned above and ranked variants. Secondly, we fine-mapped putative causal variants for each molecular phenotype by using the “Sum of Single Effects” (SuSiE) model (v 1.0) (*46*). We calculated LD correlations between all tested SNPs of a molecular phenotype from the genotype reference panel and then fine-mapped variants using the SuSiE infinitesimal effect model. The posterior probability of 0.1 was used for identifying putative causal variants and credible sets.

### Colocalization analysis between molecular phenotypes

To demonstrate whether two types of molecular phenotypes shared genetic regulatory mechanisms, we determined a set of paired molecular phenotypes that were transcribed from the same gene. We then ran the *coloc.abf* function in the coloc package(*154*), which is an Approximate Bayes Factor colocalization analysis for detecting significant genetic variants shared by two molecular phenotypes. The package computed posterior probabilities for: 1) no association with either molecular phenotype (H0); 2) association only with the first molecular phenotype (H1); 3) association only with the second molecular phenotype (H2); 3) association with both molecular phenotype but two independent signals (H3); 4) association with both molecular phenotype and shared signals (H4). Moreover, we calculated the linkage disequilibrium (LD) of two lead SNPs for a pair of shared molecular phenotypes using PLINK v1.9(*146*).

### Tissue- and breed-sharing of molQTL

*Tissue-sharing of molQTL.* To assess the cross-tissue sharing pattern of molQTL, we used Multivariate Adaptive Shrinkage in R (MashR, v0.2.57)(*109*) and METASOFT v2.0.0(*155*). For MashR, we used the z-score (slope/slope_se) of top molQTL for a gene as input. To run the *mash* model, we randomly selected 1 million molQTL-gene pairs from nominal associations being tested across all tissues by tensorQTL and obtained their z-score values. If there were missing z-score values, zero was filled and the corresponding standard error was set to 1e^6^. Local false sign rate (LFSR) was then computed by MashR and an LFSR of 0.05 was considered as the significance threshold to define whether a molQTL was active in a tissue. Pairwise Spearman’s correlation of effect size of active molQTL was calculated to evaluate tissue similarity. For METASOFT, we combined all significant molQTL across tissues and computed the z-score as described above. We estimated the m-value, which represented the posterior probability indicating whether a molQTL effect exists in a tissue, using the Markov Chain Monte Carlo (MCMC) method(*156*). The m-value threshold was set as 0.7.

*Breed-sharing eQTL analysis*. We considered the brain (Leghorn, n = 78; Red Jungle Fowl, n = 46; Ross, n = 157), spleen (Leghorn, n = 74; Cobb, n = 43) and liver (Leghorn, n = 60; Cobb, n = 47; Ross, n = 101; Rhode Island Red, n = 78), tissues as they had more than two breeds with sample size > 40. For each breed, we ran eQTL mapping independently using tensorQTL software (v1.0.4). The eQTL sharing was assessed using METASOFT v2.0.0(*155*), and MashR (v0.2.57)(*109*), as well as π1 statistic in the qvalue package(*51*, *157*). The METASOFT and MashR were run as described above, and the π1 statistic (i.e. replication rate)(*157*) was used to assess if an eQTL detected in one breed can be replicated in another breed.

### Detection of context-dependent QTL

*Sex-biased eQTL.* To identify eQTL that is significantly associated with gender, we focused on eight tissues that had at least 30 samples for each sex. In this study, we only considered conditionally independent eQTL identified above to reduce the computational burden. We fitted a linear model y = g + s + s × g + c + e, where y is phenotypic values of gene expression; g is genotype (0 for homozygous ref, 1 for heterozygous, and 2 for homozygous alt); s is sex information (0 for female and 1 for male); c is quantitative covariates including 5 genotypic PCs and 10 PEER factors as we used in eQTL mapping, while e is for the residuals. The same parameters were also used for computing the null model but excluding the s × g term. We then calculated *P* values by comparing the linear interaction model to the null model using analysis of variance. The *lm()* function in R v3.6.3 (*105*) was used for model fitting.

*Transcription factor (TF) interacting eQTL.* To detect eQTL that may interact with the expression of transcription factors, we retrieved 956 putative transcription factors from the AnimalTFDB (v3.0). As was done for sex-biased QTL detection, we only considered conditionally independent eQTL but excluded eGenes that were TF. Likewise, we fitted the same interaction model, but the interaction term was TF expression. A total of 15 quantitative covariates including 5 genotypic PCs and 10 PEER factors were also fitted in the model to control confounding factors. The significance threshold was set as FDR(*130*) corrected *P*-value < 0.01.

*Cell-type interacting eQTL*. We mapped cell-type interaction QTLs by fitting a linear model but included an interaction term implemented in the tensorQTL v1.0.4 (*152*): y = g + i + g × i + e, where y is gene expression, i is the estimated abundance of cell types, and g is genetic effects estimated from SNPs within ±1Mb of the TSS of a gene, while e is for the residuals. To control confounding factors, we also included a total of 15 quantitative covariates including 5 genotypic PCs and 10 PEER factors as described above. We defined genes that had at least one significant SNP after carrying out a multiple testing correction on eigenMT-based *P*-values(*158*) using the FDR approach(*130*). We defined the threshold of significance as FDR < 0.01.

*Breed interacting eQTL*. To demonstrate breed effects on gene regulation, we ran breed interaction eQTL mapping using the tensorQTL (v1.0.4) tool (*152*) in brain, where the sample size of each breed was > 40, including Leghorn (n=78), Red Jungle Fowl (n = 46) and Ross (n = 157). This interaction eQTL mapping fitted the same model as described for “*Cell-type interaction eQTL”* while the interaction term was breed information. The breed origins were coded as 0 for Red Jungle Fowl and 1 for Leghorn/Ross. After a multiple testing correction using the FDR approach(*130*), gene-variant pairs with FDR < 0.01 were deemed as significant.

### Estimating effect sizes of molQTL

We estimated the allelic fold change (aFC) of molQTL by employing the aFC (v0.3) Python script(*159*). The estimation was based on genotypes and molecular phenotypes (the same as molQTL mapping), as well as covariates including 5 genotypic PCs and 10 PEER factors. The 95% confidence interval of aFC was estimated by using the bootstrap method (--boot 100).

### Allele specific expression (ASE)

We conducted a haplotype-based ASE analysis through the phASER (v1.1.1) software(*160*). To exclude genomic regions with high mapping error rates, we first computed the genome mappability using GenMap (v1.3.0)(*161*) with parameters: -K 75 -E 2. The generated blacklist was fitted to the phASER (v1.1.1) tool(*160*) to phase variants from the STAR alignment BAM and VCF files with options of “--paired_end 1 --mapq 255 --baseq 10”. Using the script “phaser_expr_matrix.py”, we measured gene-level haplotypic expression for all samples with default parameters. The generated haplotypic counts files for individual samples were further aggregated by tissue by using “phaser_expr_matrix.py”. Finally, we used the “phaser_cis_var.py” script to estimate the effect size of eQTLs based on aggregated haplotypic counts. The correlation of ASE-level effect size (ASE aFC) and eQTL effect size (aFC estimated above) was computed using the Spearman’s correlation approach in R v3.6.3 (*105*).

### Replication of molQTL discovery

To assess the replication rate of molQTL discovery, we employed the π1 statistic embedded in the qvalue package(*51*, *157*). Briefly, we randomly split RNA-Seq samples into two groups - QTL discovery and validation population, when the tissue sample size was greater than 100. We ran QTL mapping in each group separately using tensorQTL (v1.0.4) (*152*) as described above. Based on replicated eQTL *P*-values, we calculated π0 value that measured the overall proportion of true null hypotheses using the *pi0est* function within the qvalue package(*157*). The π1 was thus obtained by 1- π0.

### DeepSEA model training and variant effect prediction

DeepSEA is a deep learning model initially trained for predicting variant effects in human(*50*), while in this study, it was retrained by utilizing 310 epigenomic peaks generated by the chicken FAANG consortium (*30*) and by Zhu et al. (*162*) (**Table S7**). According to sequencing type and histone marks, we categorized all 310 epigenomic peaks into groups, including ATAC, CTCF, DNaseSeq, H3K27ac, H3K23me3, H3K4me1 and H3K4me3, which were used as input for the model training using the Selene, a PyTorch-based package (*163*). Briefly, we grouped the genome into 200-bp bins and then labeled the bins according to input features. A genomic bin will be labeled 1 if half of the bin overlaps with an epigenomic peak, otherwise labeled as 0. The model was then trained based on a sequence region of 1,000 bp (i.e., input feature), where the 200-bp bin was placed at the center. We created validation and testing datasets by grouping chromosomes, specifically grouping chromosomes 8 and 9 to the training set and chromosomes 6 and 7 to the validation set. We computed the area under the receiver operating characteristic (AUROC) to evaluate the performance of the DeepSEA model. After that, we computed variant effect/score of two alleles for a given molQTL, i.e. 2 × 310 predicted chromatin variant scores, by inputting a 1000-bp sequence with the center being the Ref or Alt allele. The score is defined as the relative log fold changes of odds between predicted scores of the Ref and Alt. For each feature, SNPs with a score greater than 0.7 were identified as variants affecting the feature.

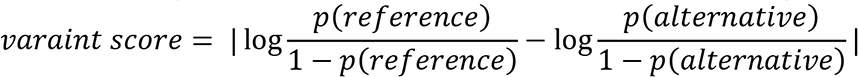

### Functional enrichment of molQTL

To understand the enrichment of molQTL in sequence ontology (i.e., SNP functional types annotated by SnpEff v5.0e) and regulatory elements (i.e., 15 chromatin states annotated in(*30*)), we employed the formula:

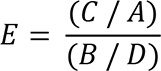

where A and D are the length of feature annotations and the total genome length, respectively. C is the length of molQTL overlapped with feature annotations, and B is the length of molQTL overlapped with the total genome length. To further uncover the regulatory mechanism of molQTL, we retrieved predicted pairs of regulatory elements-target genes from(*30*). We then overlapped them with our molQTL-regulated genes but at the same time required the molQTL to be located within regulatory elements. Moreover, we also performed the enrichment analysis of molQTL-regulated genes and HiC TAD with data retrieved from a previous study (*52*) using the SnakeHiC pipeline (https://github.com/FarmOmics/SnakeHiC).

### Integrating molQTL with GWAS results

*GWAS summary statistics.* To investigate the regulatory mechanisms underpinning complex traits in pigs, we systematically integrated the identified molQTL with GWAS from 108 complex traits of economic importance, representing five trait domains (i.e., growth and development, carcass, egg production, feed efficiency and blood biochemical index). Detailed information for each GWAS is shown in **Table S17**. To perform the integrative analysis of GWAS and molQTL, we overlapped significant GWAS loci with the 1,522,091 SNPs were tested in the molQTL mapping analysis, resulting in 1,176 GWAS loci.

*Enrichment of molQTL and trait-associated variants.* To examine whether molQTL were significantly enriched among the significant GWAS loci, we applied QTLEnrich (v2) (*32*) to quantify the enrichment degree between significant molQTL and GWAS loci.

*Transcriptome-wide association study (TWAS)*. We conducted single- and multi-tissue TWAS with S-PrediXcan(*164*) and S-MultiXcan(*165*) included in the MetaXcan (v0.6.11) family, respectively. Briefly, we trained the Nested Cross validated Elastic Net models with molecular phenotypes (i.e., PCG, lncRNA, splicing, exon, and 3a’Genes) and corresponding SNPs within the 1Mb *cis*-window of molecular phenotypes in all 28 tissues. The predictive models with cross-validated correlation ρ > 0.1 and prediction performance *P* < 0.05 were selected for subsequent analyses. Using the S-PrediXcan tool and trained models, we predicted gene-trait associations at the single-tissue level, i.e., single-tissue TWAS results. Further, using the S-MultiXcan tool, we integrated single-tissue predictions, generating the multiple-tissue TWAS results. After carrying out a multiple testing correction with the FDR approach(*130*), gene-trait associations with corrected-*P* < 0.05 were considered as significant.

*Summary-based Mendelian Randomization (SMR)*. To explore the pleiotropic association between molecular phenotypes and a complex trait, we conducted a Mendelian Randomization analysis. This was done by using the SMR software (v1.3.1) (*166*), which can utilize summary-level data from GWAS and molQTL. To correctly fit the SMR software, the molQTL data generated by tensorQTL in this study was initially converted into BESD format with options of “--fastqtl-nominal-format --make-besd”. We then ran the SMR test and carried out a multiple testing correction with the FDR approach(*130*). The gene-trait pairs with corrected *P*-value < 0.05 were selected and deemed as significant.

*Colocalization analysis*. To identify shared genetic variants between GWAS and molQTL, we conducted a colocalization analysis with fastENLOC (v2.0) (*167*). We first fine-mapped putative causal variants for each eGene by using a Bayesian multi-SNP genetic association analysis algorithm, deterministic approximation of posteriors (DAP, the current version is DAP-G, v1.0.0)(*168*, *169*). Leveraging the DAP-G (v1.0.0) (*168*, *169*) outcome, we generated a probabilistic annotation of molQTL using the “summarize_dap2enloc.pl” script. We then calculated approximate LD blocks using PLINK v1.9 (*146*) with options: --blocks no-pheno-req --blocks-max-kb 1000 --make-founders. The posterior inclusion probability (PIP) of GWAS loci was calculated for each LD block using TORUS (*170*) with the options: --load_zval -dump_pip. By integrating GWAS PIP values, we ran the final colocalization analysis with the fastENLOC (v2.0) tool (*171*) and obtained the regional colocalization probability (GRCP). The GRCP > 0.1 was defined as the threshold of significance.

### Enrichment analysis of eQTL in selective sweeps

To determine whether domestication could be acting on regulatory variants, we retrieved selective sweeps measured by locus-specific branch length (LSBL) statistics (*14*, *65*). Briefly, we first calculated FST for genomic windows with 20 consecutive SNPs between broilers (n=40) and Red Jungle Fowls (RJF, n=35) using VCFtools and also between layers (n=50) and Red Jungle Fowls (n=35). The LSBL values were further computed with the formula: LSBL = (FST(AB) + FST(AC) - FST(BC)) / 2. We deemed the top 0.1% of genomic windows ranked by LSBL values as significant. We examined whether eQTL were overrepresented in genomic windows under position selection, *i.e.,* whether genomic windows with at least one eQTL had higher LSBL values than the background, which has an equivalent number of windows as those of eQTL.

### Comparative analysis of gene expression

To comparatively analyze gene expression across species, we collected gene expression and regulation data from multiple sources. Specifically, we obtained data for 15,044 samples from the Human GTEx web portal (v8) available at https://gtexportal.org. Additionally, we gathered gene expression data for 7,095 pig samples and 8,742 cattle samples from the FarmGTEx resource accessible at https://www.farmgtex.org/. Furthermore, as part of this study, we included gene expression and regulation data for 7,015 chicken samples. In this study, we focused on protein-coding genes based on the annotation of the Ensembl (v102), and we considered the genes with TPM > 0.1 as expressed. Specifically, we grouped chicken genes into “1-1-1-1 orthologous gene” (1-1 orthologous across species, n = 10600), “complex orthologous genes” (“1 to many”, “many to 1” and “many to many”, n=3644), “no homology” (without any homologous counterpart in mammals, n=2535). In total, 9 tissues in common across species (*i.e.,* adipose, blood, hypothalamus, liver, lung, muscle, ovary, pituitary, and testis) were included to conduct a comparative analysis of gene expression.

Gene expression (TPM matrix) retrieved for each was normalized using Seurat (v4.3.0) (*110*) to decrease the bias introduced by dynamic data across species. We evaluated transcriptome outcomes by counting the number of reads (reads = TPM × the length of genes (bp)) in each tissue. Samples were by performing dimensionality reduction on the normalized expression data (including 10,600 1-1-1-1 orthologous genes) with the *t*-SNE approach (*117*). Moreover, we explored the conservation of cis-heritability (*h^2^*) and effect size (aFC) of lead eVariants across species. To do so, we selected 1-1-1-1 orthologous genes (n=5,384 for *h^2^*) and eGenes related eVariants (n=5,513 for aFC) that are in common across species, and grouped genes into conserved eGenes (that have at least 1 homology with mammals) and chicken-specific eGenes (that didn’t have homology with any mammal species).

### Cross-species TWAS comparison

We performed comparative analyses of single-tissue TWAS results of 108 traits in chickens with 9,112, 1,032 and 6,480 single-tissue TWAS results in three mammalian species (*i.e.*, pigs(*47*), cattle(*48*) and humans(*32*)), representing 268, 43 and 135 complex traits, respectively. Within a shared tissue between two species, we computed Pearson’s correlations between any pair of traits on the basis of z score (beta / standard error) estimated from one-to-one orthologous genes between two corresponding species. To define the threshold of significance, we carried out permutation analysis by randomly calculating Pearson’s correlations between any two of all single-tissue TWAS 1000,000 times. The within-species correlations were then excluded, resulting in 609,861 Pearson’s correlations and corresponding *P* values. We set the cutoff of significance as top 0.1% of permuted -log10*P*, corresponding to the *P* value of 9.11 × 10^-3^. We thus conducted cross-species meta-TWAS analysis by combining TWAS results from different species based on orthologous genes. For meta-TWAS analysis, we applied a sample-size weighting (SSW) strategy (*172*) by calculating *ZTWAS* as follows:

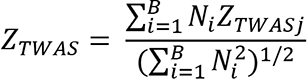

where *ZTWASj* is the z-score for *j*th gene in TWAS analysis, *i* is the species, *i.e.*, chicken, humans, pigs, and cattle, *Ni* is the number of individuals for *i*th species in TWAS, *B* is the number of species in metaTWAS. The effective sample size is 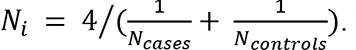. To obtain the significance level, we calculated *P* values for each gene based on a Chi-squared distribution of z-scores (df=1) calculated before. After a multiple testing correction with the FDR method (*130*) by replacing original *P* value (TWAS) with *P* value (meta-TWAS) of orthologous genes, the threshold of significance was defined as FDR < 0.05.

## Supplementary Materials

Materials and Methods Figs. S1 to S32

Tables S1 to S22

